# Animal-vectored nutrient flows across resource gradients influence the nature of local and meta-ecosystem functioning

**DOI:** 10.1101/2023.03.03.530982

**Authors:** Matteo Rizzuto, Shawn J. Leroux, Oswald J. Schmitz, Eric Vander Wal, Yolanda F. Wiersma, Travis R. Heckford

## Abstract

Organisms moving across landscapes connect ecosystems in space and time, mediating nutrient, energy, and biomass exchanges. Meta-ecosystem ecology offers a framework to study how these flows affect ecosystem functions in space and time. However, meta-ecosystem models often represent consumer movement as diffusion along gradients of resources. Crucially, this assumes that consumer movement connects the same trophic compartments among patches of the same ecosystem. Yet, empirical evidence shows that organisms move across different ecosystems and connect diverse trophic compartments in diffusive and non-diffusive ways. Here, we derive a two-patch meta-ecosystem model that accounts for both types of organismal movement, and we investigate their influences on local and meta-ecosystem functions. We integrate two novel approaches in this classic meta-ecosystem model: a dispersers’ pool to capture the fraction of moving organisms and time scales separation to partition local and regional dynamics. We show that non-diffusive consumer movement increases landscape heterogeneity while diffusive consumer movement enhances source-sink dynamics. Local ecosystem differences driven by consumer movement type are less prevalent at meta-ecosystem extents. Thus, movement type is essential for predicting local ecosystem dynamics. Our results support recent calls to explicitly consider the role of consumers in shaping and maintaining ecosystem functions in space and time.

## 1 Introduction

Ecosystems are intrinsically open systems connected by exchanges of different currencies (e.g., energy, matter, information; Loreau, Mouquet, and Holt 2003; Marleau, Peller, et al. 2020; Little et al. 2022). Abiotic and biotic vectors—for instance, rivers or migratory animals—drive these exchanges and thus influence ecosystem functioning at multiple spatial extents (Gravel, Guichard, et al. 2010; Schiesari et al. 2019). While the influences of abiotic vectors on ecosystem dynamics have been extensively studied (Gravel, Guichard, et al. 2010; Gravel, Mouquet, et al. 2010; Gounand, Mouquet, et al. 2014; Loreau, Daufresne, et al. 2013), less is known about how flows mediated by biotic vectors affect ecosystem processes and functions (Gounand, Harvey, et al. 2018; but see Subalusky, Dutton, Njoroge, et al. 2018; Schmitz, Wilmers, et al. 2018). Evidence from both the fossil record and present-day events shows that biotic drivers of ecosystem flows affect ecosystem functions at extents ranging from local to continental (Bauer and Hoye 2014; Schmitz, Raymond, et al. 2014; Doughty 2017; MacSween, Leroux, and Oakes 2019). However, organism-driven exchanges have diminished over time as humankind began modifying the biosphere (from the late Quaternary onwards; Doughty et al. 2016). Mathematical models can play an increasingly key role in clarifying and predicting causes and consequences of human-driven changes to abiotic and biotic flows connecting ecosystems over space and time (McCann et al. 2021). Most models and theory on organismal movement, however, focus on patterns at the population and community level (reviewed in Bauer and Hoye 2014), without addressing their ecosystem effects. The ensuing conceptual and practical gap hampers the study of how organismal movement impacts ecosystem functions, in particular biogeochemical cycling, at local and regional extents (i.e., zoogeochemistry; sensu Schmitz, Wilmers, et al. 2018).

Organisms can move large quantities of nutrients across ecosystem borders, influencing donor and recipient ecosystems (Earl and Zollner 2014, 2017; Hobbie and Villóeger 2015). For instance, brown bear (*Ursus arctos*) consume migratory Pacific salmon (*Oncorhynchus* spp.) and through their against-gradient movement transfer ocean-derived nutrients to the riparian forests surrounding their spawning streams in north-western North America, increasing primary productivity at the forest stand level (Francis, Schindler, and Moore 2006; Helfield and Naiman 2001). Similar animal-mediated subsidies drawn from ocean depths can benefit productivity once released in ocean surface waters (Roman and McCarthy 2010). In terrestrial ecosystems, large herbivores are well-known vectors of nutrient exchanges (Seagle 2003; Frank 2008; Abbas et al. 2012). In Kenya, for instance, hippopotamuses (*Hippopotamus amphibius*) and other large herbivores transfer large quantities of nitrogen from grasslands to the Mara River (Subalusky, Dutton, Rosi-Marshall, et al. 2015; Subalusky, Dutton, Njoroge, et al. 2018). Analogous flows act in circumpolar (Jefferies, Rockwell, and Abraham 2004), boreal (Seagle 2003; Bump et al. 2009), and temperate (Abbas et al. 2012) biomes, mediated by local and migratory animals. These organismal nutrient subsidies affect fundamental ecosystem processes, such as primary productivity (Helfield and Naiman 2001; Subalusky, Dutton, Njoroge, et al. 2018) and nutrient cycling (Abbas et al. 2012; Bump et al. 2009). What is more, these subsidies often relate to the ratio of subsidy biomass to local resource biomass. Consequently, the shape of biomass pyramids (sensu McCauley et al. 2018) in donor and recipient ecosystems may impact the outcome of subsidies. Human activities stand to continue to modify these widespread organismal-determined pathways across ecosystems as we progress in the Anthropocene.

Meta-ecosystem theory—which studies how flows of energy, matter, and organisms connect ecosystems in space and time (Loreau, Mouquet, and Holt 2003; Gounand, Harvey, et al. 2018)— is a useful framework to place the empirical evidence for organismal pathways across ecosystems in the broader context of an interconnected biosphere. Organism-mediated flows of nutrients bridge ecosystem processes and functions, changing the structure and properties of local and meta-ecosystems (Leroux and Loreau 2012; Massol, Altermatt, et al. 2017; McCauley et al. 2018). Regional, consumer-mediated flows of nutrients across ecosystems can have cascading effects on local biomass recycling processes, primary and secondary production, and ecosystem stability (Marleau, Guichard, Mallard, et al. 2010; Marleau, Guichard, and Loreau 2014). For instance, these flows can sustain autotroph populations in ecosystems with low nutrient availability and allow them to thrive (Gravel, Guichard, et al. 2010; Subalusky, Dutton, Rosi-Marshall, et al. 2015). Organismal movement can further benefit primary producers by redistributing nutrients over regional spatial extents, alleviating the effects of excessive nutrient availability (i.e., ameliorating the Paradox of Enrichment; Gounand, Mouquet, et al. 2014). As well, the movement of organisms allows trophic interactions in one ecosystem to influence those of adjacent ones, producing spatial trophic cascades whereby apical trophic compartments from donor ecosystems influence basal compartments in recipient ones (sensu Knight et al. 2005; Massol, Altermatt, et al. 2017). Through modifications of regional source-sink dynamics, biotic subsidies can thus shift the distribution of biomass and abiotic nutrients stocks in space and time (Gravel, Guichard, et al. 2010; Gounand, Mouquet, et al. 2014).

Recently, meta-ecosystem theory has been criticized for having weak linkages to empirical research (Gounand, Harvey, et al. 2018; Schiesari et al. 2019; Peller, Guichard, and Altermatt 2023). We surmise that one of the reasons for this is that most meta-ecosystem theory treats organismal movements as diffusive flows along environmental gradients—e.g., from high to low resource availability (Gravel, Guichard, et al. 2010; Marleau, Guichard, Mallard, et al. 2010; but see Leroux and Loreau 2012). Assuming diffusive movement of organisms implicitly assumes that it occurs among patches within the *same* ecosystem, because the *same* organisms are moving to or from the *same* compartment from one patch to another (sensu Massol, Altermatt, et al. 2017). That is, diffusion-like movement deprives consumers of agency (Gounand, Harvey, et al. 2018; Little et al. 2022) in navigating the landscape. However, organisms routinely cross the boundaries of different ecosystem types in against-gradient or gradient-neutral ways (i.e., from low to high resource availability or in the absence of a gradient, respectively; Gounand, Harvey, et al. 2018). Through these movements, organism biomass is often converted from one compartment to another (Gounand, Harvey, et al. 2018), for instance, via ontogenic niche shifts (e.g., juvenile insectivorous salmons in rivers evolve to adult piscivore salmons in the ocean; Ebel et al. 2015) or senescence (e.g., leaves falling into freshwater bodies). Merely assuming diffusive organismal movement also overlooks evidence—from wildlife and behavioral ecology—of the pervasive fitness trade-offs that drive organismal movement (Hugie and Dill 1994; Nathan et al. 2008). As organisms move over landscapes, short-term (e.g., avoiding predation, competition, starvation) and long-term (e.g., growth, reproduction) needs influence the processes of searching, entering, and foraging in new patches (Nathan et al. 2008; Gounand, Harvey, et al. 2018; Little et al. 2022). Furthermore, the matrix surrounding local ecosystem patches also matters, as organisms traversing it do not contribute to ecosystem dynamics while they remain in the matrix or if they succumb to mortality while there (Figure 1a; Weisser and Hassell 1996).

**Figure 1:**
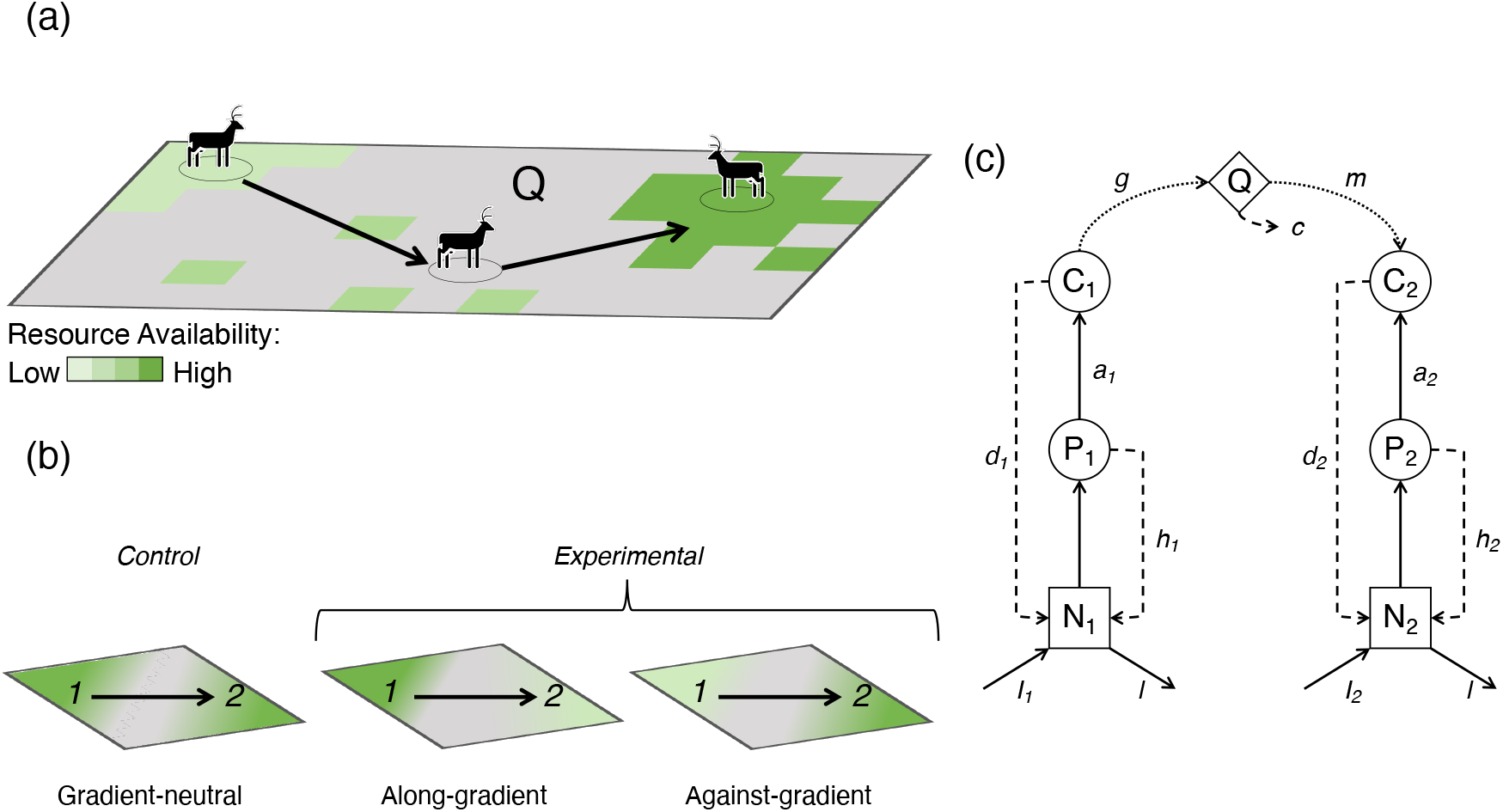
Visual concepts for the (a) rationale, (b) analyses, and (c) structure of the model. **Panel (a)**: Organisms, i.e., consumers, move among ecosystems over a gradient of resource availability (green-shaded polygons) by traveling across the matrix (grey areas, 𝒬), thus connecting and potentially influencing ecosystem functions at local and regional spatial scales. **Panel (b)**: We consider three scenarios of consumer movement over gradients of nutrient availability (*I*_*i*_; Table 1), here shown in green. Organisms move from ecosystem 1 (donor) to ecosystem 2 (recipient) in all scenarios (black arrow). The grey-shaded area in the middle of each scenario represents the unsuitable matrix 𝒬 between donor and recipient ecosystems. In the control scenario, on the left, the two ecosystems have equal nutrient availability (*I*_1_ = *I*_2_) and consumer movement is gradient-neutral. In the first experimental scenario, center, the donor ecosystem has higher nutrient availability than the recipient ecosystem (*I*_1_ >> *I*_2_) and organisms move along gradient. In the second experimental scenario, right, the recipient ecosystem has higher nutrient availability than the donor ecosystem (*I*_1_ << *I*_2_) and organisms move against gradient. **Panel (c)**: Diagram of the model, showing two local ecosystems connected by unidirectional movement (dotted lines) through a dispersers’ pool 𝒬. Boxes and circles represent nutrient stock, primary producers, and consumers trophic compartments, respectively. Solid arrows connecting trophic compartments represent trophic interactions and dashed arrows represent recycling pathways. See Table 1 for definitions of the variables and parameters in the model.

Here, we aim to (i)) investigate the influence of consumer movement on ecosystem functions at local and regional extents and (ii) enhance the ecological realism of meta-ecosystem theory, by integrating different types of organismal movement over resource availability gradients into a classic meta-ecosystem model.We focus on three types of consumer movement with respect to resource availability gradients: gradient-neutral, along-gradient, and against-gradient movement. Our model combines a two-patch meta-ecosystem with an intermediate compartment for dispersing consumers (henceforth, dispersers’ pool; Weisser and Hassell 1996). While not spatially explicit, including this disperses’ pool provides a first approximation of the inhospitable matrix that consumers traverse when moving between ecosystems, echoing the “patch-matrix mosaic” model frequently used in landscape ecology to describe real-world landscapes (Castilla et al. 2009; Wu 2013). Following other ecosystem models (Ludwig, Jones, and Holling 1978; Menge, Pacala, and Hedin 2009; Menge, Hedin, and Pacala 2012), we use time scales separation to account for the different scales of movement in our meta-ecosystem and account for the complex temporal dynamics of our meta-ecosystem model (Otto and Day 2011). We investigate how these different biotic movement types influence local and meta-ecosystem functions, e.g., biomass and nutrient stock accumulation, nutrient flux, and primary and secondary productivity. We expect that different types of consumer movement will influence local and regional ecosystem functions, albeit not necessarily in the same direction or with the same magnitude (Figure 1b; Marleau, Guichard, Mallard, et al. 2010).

## 2 Ecosystem Model

### 2.1 Model Derivation

We derive a meta-ecosystem model comprising two ecosystems (eqs. (1a) to (1g); Figure 1c), connected by movement of consumers mediated through a dispersers’ pool (sensu Weisser and Hassell 1996). Table 1 summarizes the model’s state variable and parameters, their units of measurement, and any constraints on their values. Each ecosystem contains a food web composed of three trophic compartments whose biomass is measured in grams (g): inorganic nutrients (*N*_*i*_), primary producers (*P*_*i*_), and consumers (*C*_*i*_)—where *i* ∈ (1, 2) indicates the two ecosystems. The meta-ecosystem obeys mass balance constraints, and both ecosystems are open at the basal level through constant inputs (*I*_*i*_) and output (*l*) of inorganic nutrients. Biotic compartments lose biomass at rate *h*_*i*_ for producers and at rate *d*_*i*_ for consumers. Following Gravel, Guichard, et al. (2010), we assume all biomass lost from higher trophic levels (*P*_*i*_, *C*_*i*_) re-enters each ecosystem at the basal level (*N*_*i*_). Producers acquire nutrients from the basal level of each ecosystem at rate *u*_*i*_. We assume a linear functional response (i.e., Type I; Holling 1959; Jonsson 2017): consumers uptake producer biomass at rate *a*_*i*_ and convert it to new consumer biomass with efficiency *ϵ*_*i*_. Using a Type I functional response presents several advantages (Jonsson 2017): in addition to its simple formulation, which allows us to maintain mathematically tractability of the model, it has been extensively studied since first proposed and widely used in previous meta-ecosystem models (see review in Guichard and Marleau 2021); thus allowing easier and more direct comparison of our results to earlier studies.

**Table 1:**
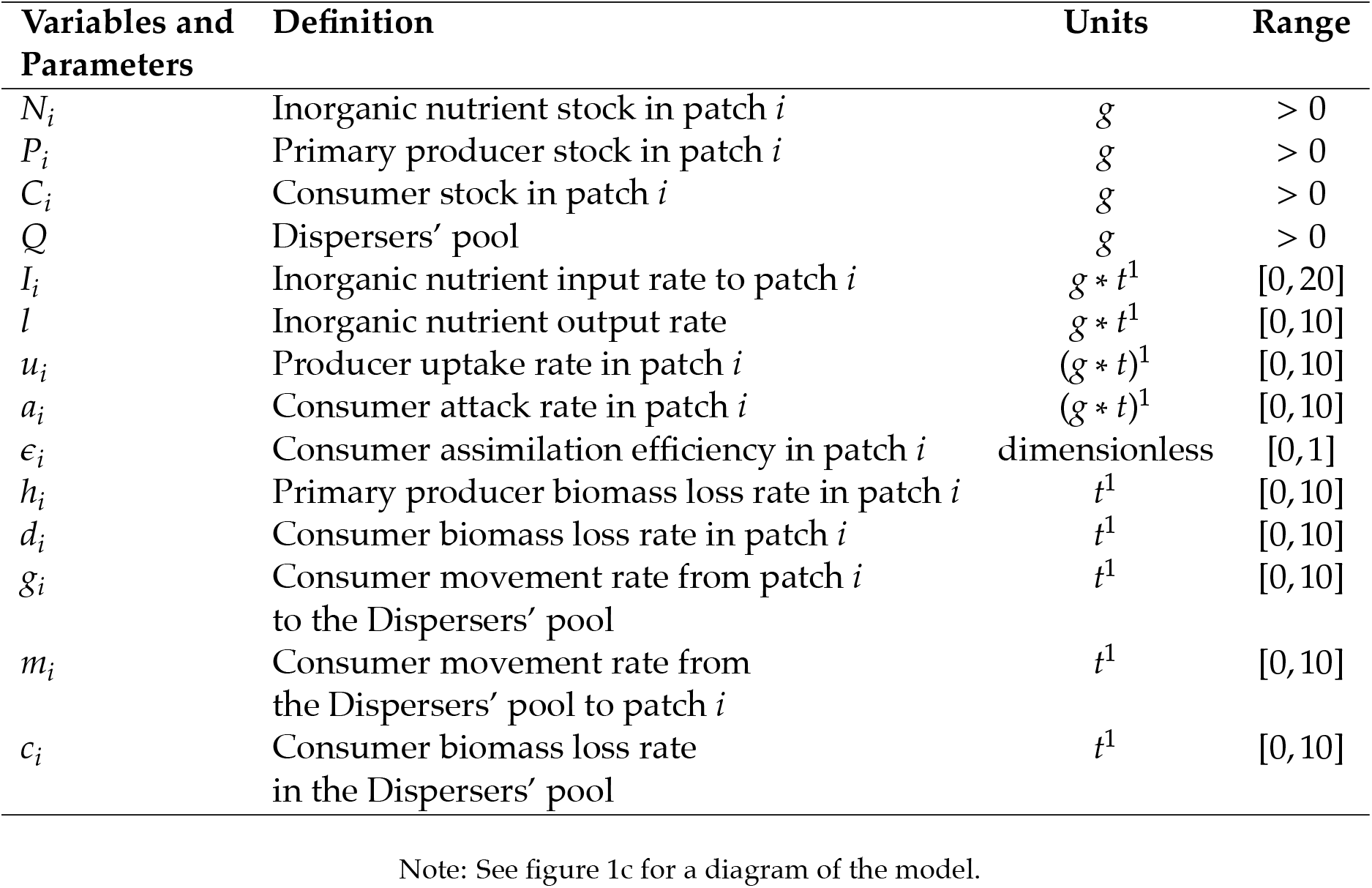
The model state variables and parameters, definitions, units of measurements, and range of values.

| Variables and Parameters | Definition | Units | Range |
| --- | --- | --- | --- |
| $N_i$ | Inorganic nutrient stock in patch $i$ | $g$ | $> 0$ |
| $P_i$ | Primary producer stock in patch $i$ | $g$ | $> 0$ |
| $C_i$ | Consumer stock in patch $i$ | $g$ | $> 0$ |
| $Q$ | Dispersers' pool | $g$ | $> 0$ |
| $I_i$ | Inorganic nutrient input rate to patch $i$ | $g * t^1$ | $[0, 20]$ |
| $l$ | Inorganic nutrient output rate | $g * t^1$ | $[0, 10]$ |
| $u_i$ | Producer uptake rate in patch $i$ | $(g * t)^1$ | $[0, 10]$ |
| $a_i$ | Consumer attack rate in patch $i$ | $(g * t)^1$ | $[0, 10]$ |
| $\epsilon_i$ | Consumer assimilation efficiency in patch $i$ | dimensionless | $[0, 1]$ |
| $h_i$ | Primary producer biomass loss rate in patch $i$ | $t^1$ | $[0, 10]$ |
| $d_i$ | Consumer biomass loss rate in patch $i$ | $t^1$ | $[0, 10]$ |
| $g_i$ | Consumer movement rate from patch $i$ to the Dispersers' pool | $t^1$ | $[0, 10]$ |
| $m_i$ | Consumer movement rate from the Dispersers' pool to patch $i$ | $t^1$ | $[0, 10]$ |
| $c_i$ | Consumer biomass loss rate in the Dispersers' pool | $t^1$ | $[0, 10]$ |

Note: See figure 1c for a diagram of the model.

We model consumer movement via a matrix between the two ecosystems using a dispersers’ pool 𝒬 (eq. (1g); Weisser and Hassell 1996) to represent that matrix. Consumers move from the donor ecosystem 1 to 𝒬 at rate *g*, and then move from 𝒬 to the recipient ecosystem 2 at rate *m* (Figure 1c). Consumer in 𝒬 do not produce new biomass and extant biomass in 𝒬 can be lost from the system at rate *c*, reflecting mortality in the matrix during dispersal. The following set of ordinary differential equations describes the dynamics of the meta-ecosystem:

Ecosystem 1 (donor):

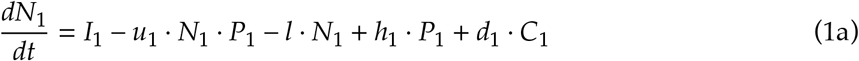

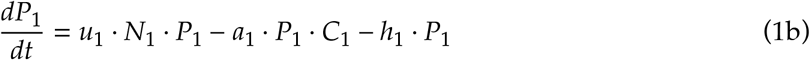

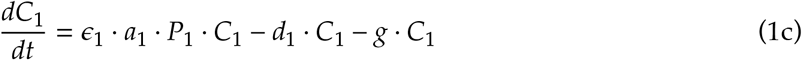

Ecosystem 2 (recipient):

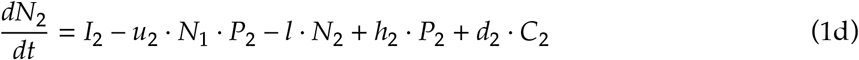

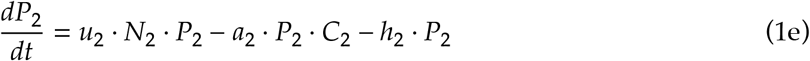

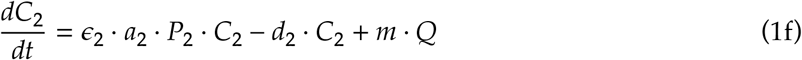

Dispersers’ pool:

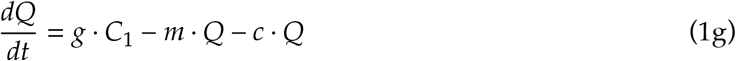

We simulate gradient-neutral, along-, and against-gradient consumer movement by keeping the structure of the model (eqs. (1a) to (1g)) constant and varying the relative values of inorganic nutrients inputs (*I*_*i*_) into each ecosystem. In turn, this varies the nutrient availability for primary producers and hence forage availability for consumers. For gradient-neutral consumer movement, we set *I*_1_ = *I*_2_, so that consumers effectively move between homogeneous ecosystem. For along-gradient consumer movement, we set *I*_1_ >> *I*_2_. Finally, for against-gradient consumer movement, we set *I*_1_ << *I*_2_.

We use time scales separation (TSS) to analyze meta-ecosystem dynamics (Otto and Day 2011). TSS helps to disentangle complex temporal dynamics in systems where multiple processes overlap in time and has a long history of application to ecosystem studies (Ludwig, Jones, and Holling 1978; Menge, Pacala, and Hedin 2009; Menge, Hedin, and Pacala 2012). Briefly, ecosystem processes can happen over long (*slow*) or short (*fast*) time scales (sensu Carpenter and Turner 2000). By assuming that *slow* process are invariant over the short scales of *fast* processes, TSS allows for finding quasi-equilibria for *fast* processes. These can then be used to find equilibria for *slow* processes (Otto and Day 2011). Here, we focus on two processes influencing the dynamics of our model: biomass production and consumer movement. In terrestrial systems, biomass production is a *slow* process taking place over months and years, whereas consumer movement is a *fast* process happening over hours or days. Consequently, we assume constant biomass production in either ecosystems while consumers move from ecosystem 1 to 𝒬 and then to ecosystem 2. Solving eq. (1g) first, we find the quasi-equilibrium 𝒬*:

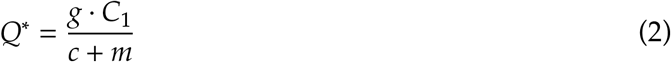

that we then substitute in eq. (1f) to solve for the *slow* processes and find equilibrium values for each trophic compartment using Mathematica (v. 13.0; Wolfram Research Inc. 2020).

### 2.2 Numerical Analyses

Our model has a single equilibrium where all state variables are positive and thus biologically meaningful (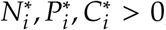; see Appendix A.1). All our analyses use this equilibrium and we run all analyses in R (v. 4.2.0; R Core Team 2022). We focus on changes caused by consumer movement on key ecosystem processes and on its interactions with meta-ecosystem biomass distribution. For both control and experimental scenarios, before running our model, we manually set the values for parameters *I*_1_, *I*_2_, and *l* as described above. For all other parameters in the model, we use 10 000 random sets of parameter values drawn from a uniform distribution using a Latin hypercube sampling design (LHS; Leroux and Schmitz 2015) and function randomLHS in the lhs R package (Carnell 2022). Following White et al. (2014), we do not conduct frequentist statistical analyses on our datasets as they arise from running large sample size numerical simulations.

#### 2.2.1 Effects of consumer movement on ecosystem processes

We assess the effects of consumer movement on ecosystem processes by estimating values of three ecosystem functions at local and regional extents (Table 2). These are nutrient stock and biomass accumulation, nutrient flux, and trophic compartment productivity. At the local ecosystem scale, we use 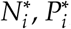, and 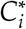 as estimates of equilibrium nutrient stock and biomass accumulation (g). We estimate local ecosystem’s primary and secondary productivity at equilibrium using the functional responses of primary producers and consumers, respectively. We estimate nutrient flux using the equilibrium loss terms of primary producers and consumers. At the meta-ecosystem scale, we sum local values of these three functions of interest. Note that meta-ecosystem nutrient flux accounts for consumer biomass lost while moving across ecosystems, i.e., parameter *c* in eq. (1g), as it does not re-enter local recycling pathways. We use the *log*_10_-transformed response ratio (*LRR*_*j*_, where 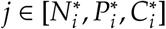) of a function’s experimental and control values to assess whether different types of consumer movement cause an increase (*LRR*_*j*_ > 0), decrease (*LRR*_*j*_ < 0), or have no effect (*LRR*_*j*_ = 0) on a given ecosystem function. Below, we report median *LRR*_*j*_ effect sizes and their ranges (see Figure A.1 in Appendix A.3 for a primer on interpreting *LRR* graphs).

**Table 2:**
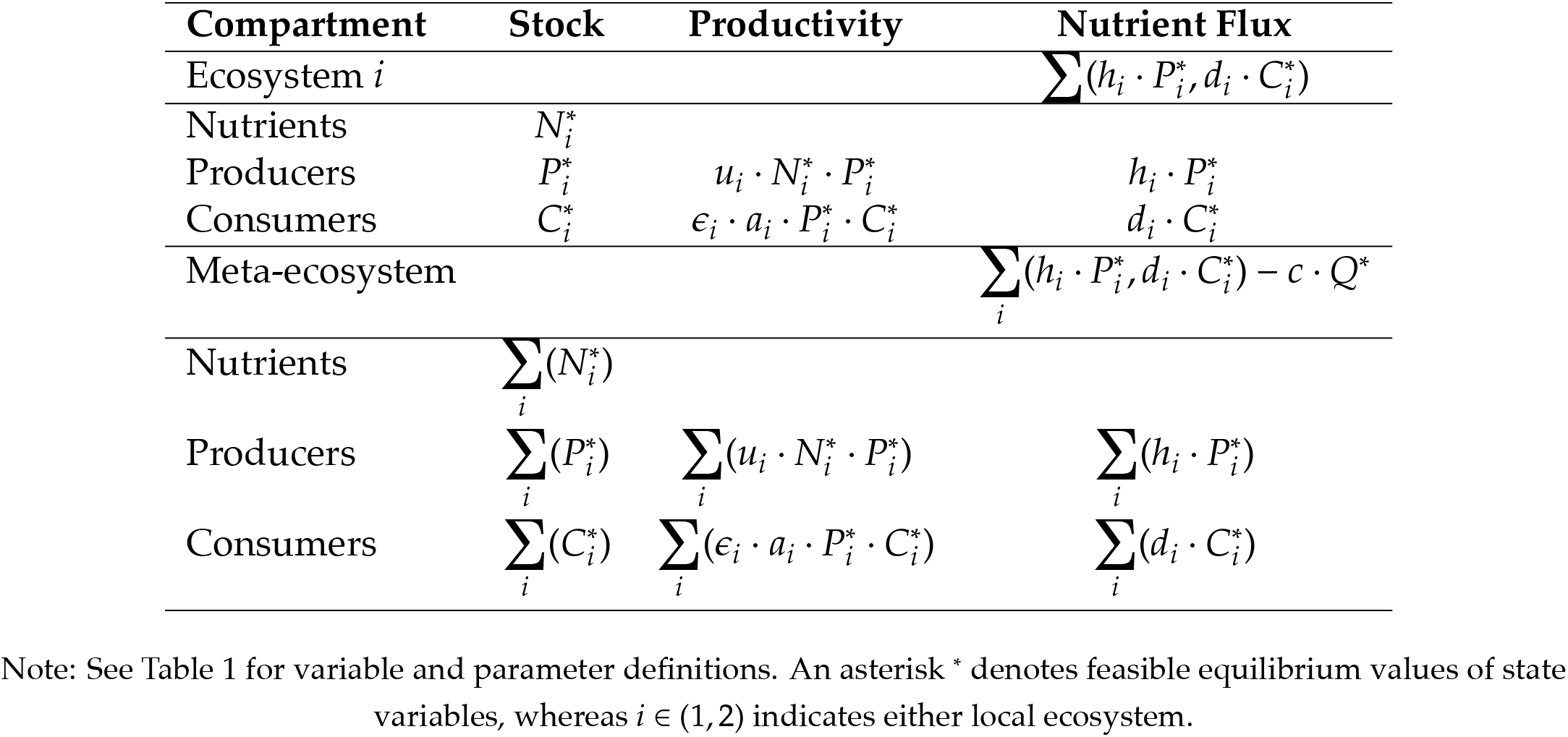
Formulae used to calculate local and meta-ecosystem functions.

| Compartment | Stock | Productivity | Nutrient Flux |
| --- | --- | --- | --- |
| Ecosystem $i$ | | | $\sum (h_i \cdot P_i^*, d_i \cdot C_i^*)$ |
| Nutrients | $N_i^*$ | | |
| Producers | $P_i^*$ | $u_i \cdot N_i^* \cdot P_i^*$ | $h_i \cdot P_i^*$ |
| Consumers | $C_i^*$ | $\epsilon_i \cdot a_i \cdot P_i^* \cdot C_i^*$ | $d_i \cdot C_i^*$ |
| Meta-ecosystem | | | $\sum_i (h_i \cdot P_i^*, d_i \cdot C_i^*) - c \cdot Q^*$ |
| Nutrients | $\sum_i (N_i^*)$ | | |
| Producers | $\sum_i (P_i^*)$ | $\sum_i (u_i \cdot N_i^* \cdot P_i^*)$ | $\sum_i (h_i \cdot P_i^*)$ |
| Consumers | $\sum_i (C_i^*)$ | $\sum_i (\epsilon_i \cdot a_i \cdot P_i^* \cdot C_i^*)$ | $\sum_i (d_i \cdot C_i^*)$ |

Note: See Table 1 for variable and parameter definitions. An asterisk \* denotes feasible equilibrium values of state variables, whereas $i \in (1, 2)$ indicates either local ecosystem.

#### 2.2.2 Assessing the effects of biomass distribution

Recent empirical (Trebilco, Baum, et al. 2013; Trebilco, Dulvy, et al. 2016) and theoretical (McCauley et al. 2018)work provides evidence show that biomass distribution can impact local ecosystem dynamics. Hence, we investigate how changes in the distribution of biomass in the donor, recipient, and meta-ecosystem enabled by consumer movement impact our results. We use the Consumer to Resource (*C*:*R*) biomass ratio calculated in each local ecosystem and for the meta-ecosystem to quantify changes in the distribution of biomass in the meta-ecosystem, following consumer movement (McCauley et al. 2018). As the focus of our model is the movement of apical consumers, we use the equilibrium biomass values of consumers 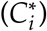 and of primary producers 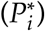 as the numerator and denominator in *C*:*R*, respectively. Values of *C*:*R* < 1 identify ecosystems with a bottom-heavy biomass pyramid, whereas *C*:*R* > 1 identifies inverted and top-heavy biomass pyramids (McCauley et al. 2018). Below, we report median *C*:*R* effect sizes and their ranges.

#### 2.2.3 Global Sensitivity Analysis

We identified the parameters with the largest influence on our model results and predictions by performing a Global Sensitivity Analysis (GSA; sensu Harper, Stella, and Fremier 2011; Bellmore et al. 2014). Briefly, we used a random forest algorithm to calculate the residual sum of squared errors (SSE) for each parameter. This parameter-specific residual SSE can then be converted into a metric of relative importance of each parameter by normalizing it against the total SSE (Liaw and Wiener 2002; Bellmore et al. 2014). We ran our GSA using function randomForest in the R package of the same name (Liaw and Wiener 2002), the 10 000 LHS-generated parameter sets described above, and the predicted equilibrium values of each state variable in the model. See section Sensitivity Analysis in the Supplementary Code document in the Data and Code repository for further details (Rizzuto et al. 2021).

## 3 Results

Consumer movement across local ecosystem borders establishes a spatial trophic cascade (sensu Knight et al. 2005; Monk and Schmitz 2022) that influences local and meta-ecosystem functions in all nutrient availability scenarios (Figures 2 to 4 and Figure A.2). Differences in the direction of consumer movement—either along- or against-gradient of nutrient availability—alter the effects and strength of this spatial trophic cascade. As well, consumer movement changes the distribution of biomass in our system, which has some ramifications for local and meta-ecosystem functioning and stability (Figures A.3 and A.6).

**Figure 2:**
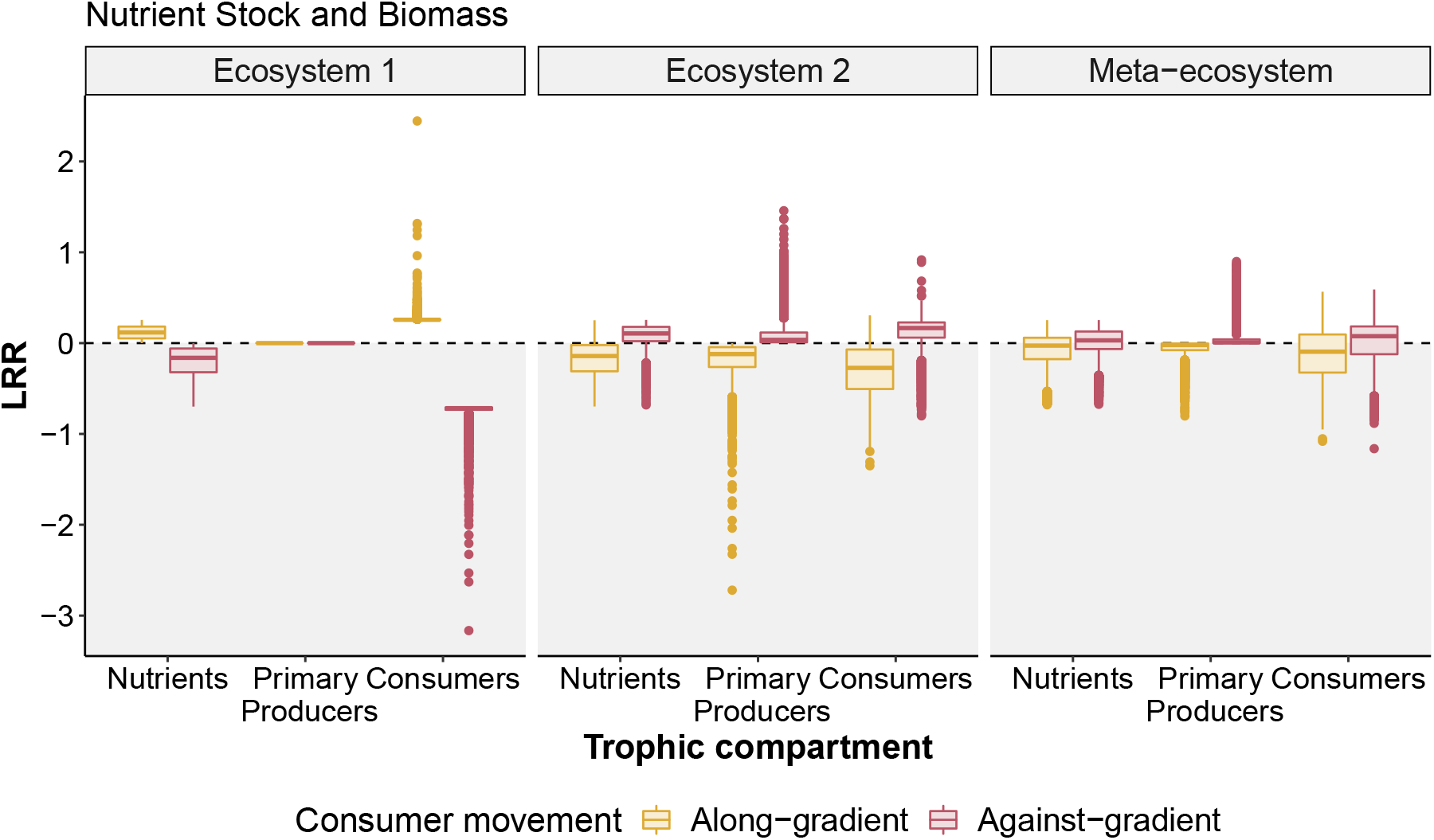
Along-(yellow) and against-gradient (red) consumer movement from ecosystem 1 to ecosystem 2 alters biomass distribution and nutrient stocks at local and meta-ecosystem extents. Along-gradient, diffusive movement (yellow) of consumers mediates higher median consumer biomass accumulation and nutrient stocks in ecosystem 1, and a corresponding reduction in ecosystem 2, compared with the control scenario. Primary producers biomass accumulation appears unaffected in ecosystem 1 and slightly reduced in ecosystem 2. This source-sink dynamic maintains meta-ecosystem nutrient stocks and biomass accumulation at levels comparable to the control scenario. Against-gradient, non-diffusive consumer movement (red) conversely mediates a spatial trophic cascade whereby ecosystem 2 shows higher median levels and ecosystem 1 has lower median values of consumer biomass accumulation and inorganic nutrient stock compared to the control, respectively. The effects of this spatial trophic cascade scale up to the meta-ecosystem, which also shows increased median consumer biomass accumulation and nutrient stocks than the control. The response ratio (*LRR*) on the ordinate captures change in local and meta-ecosystem functions in the experimental scenarios (*I*_1_ >> *I*_2_ and *I*_1_ << *I*_2_) compared to the control (*I*_1_ = *I*_2_); note its *log*_10_ scale. In each panel, thick lines inside the boxes are median values, the lower (upper) hinge is the 25% (75%) quartile, and the lower (upper) whisker extends from the hinge to the smallest (largest) values no further than 1.5 × interquartile range. Filled dots beyond the whiskers are outliers. The area above the dashed black line (white background) shows an *LRR* increase with consumer movement, while the area below it (grey background) shows a reduction.

**Figure 3:**
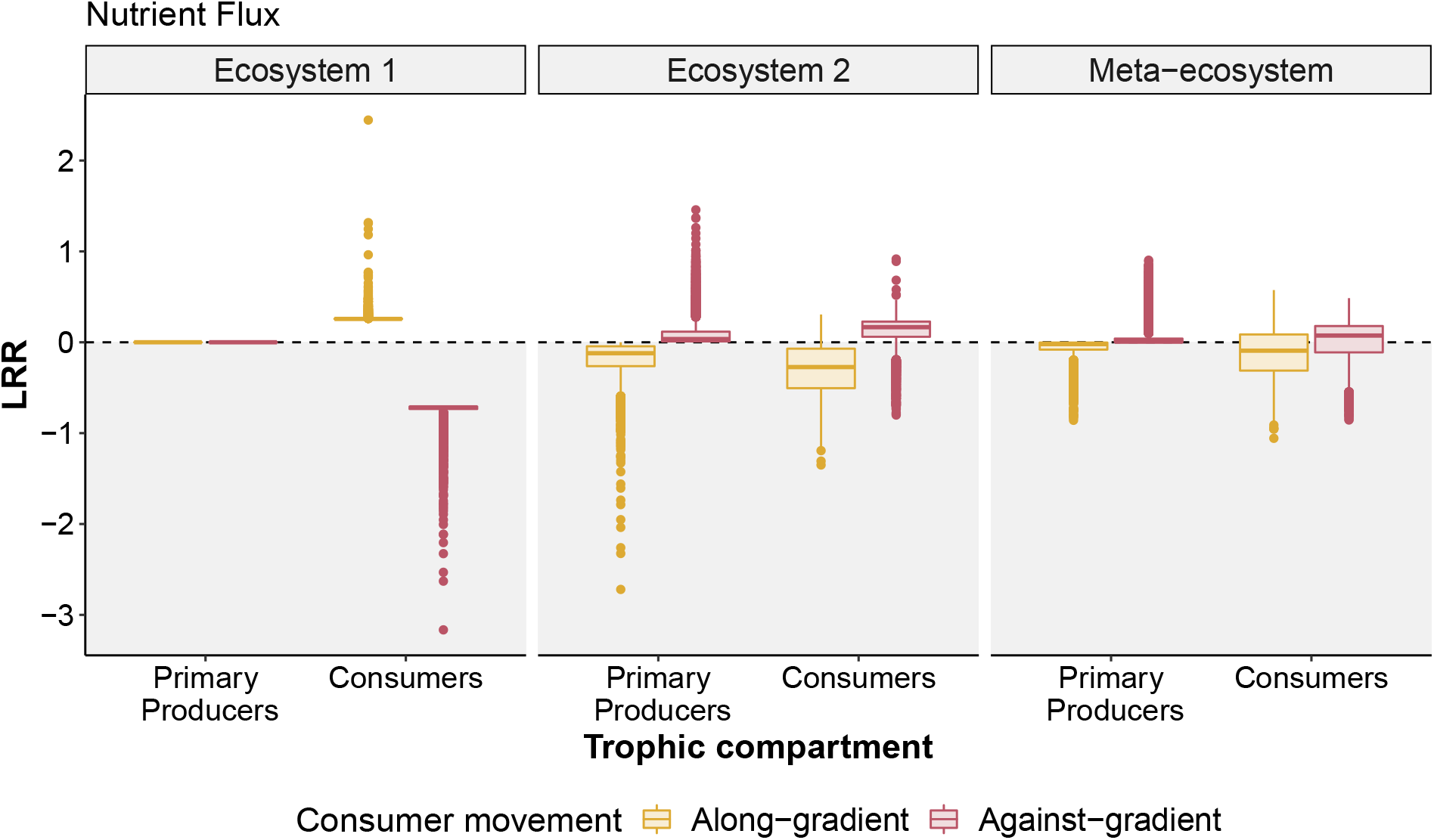
Local and meta-ecosystem nutrient flux changes with along-(yellow) and against-gradient (red) consumer movement from ecosystem 1 to ecosystem 2. Along-gradient consumer movement (yellow) appears to increase median consumer nutrient flux in ecosystem 1, leaving primary producers-mediated flux unaffected. In ecosystem 2, along-gradient consumer movement reduces median nutrient flux of both primary producers and consumers, maintaining meta-ecosystem flux at levels comparable to those of the control scenario. For against-gradient consumer movement (red), the effects are opposite: ecosystem 2 shows higher median levels of consumer-mediated nutrient flux compared to the control, whereas we see a strong reduction in ecosystem 1. Primary producers-mediated flux appears largely unaffected in either local ecosystem, in this scenario. At the meta-ecosystem extent, consumer nutrient flux appears marginally increased compared to the control, when against-gradient movement connects the local ecosystems. All specifications as in Figure 2.

**Figure 4:**
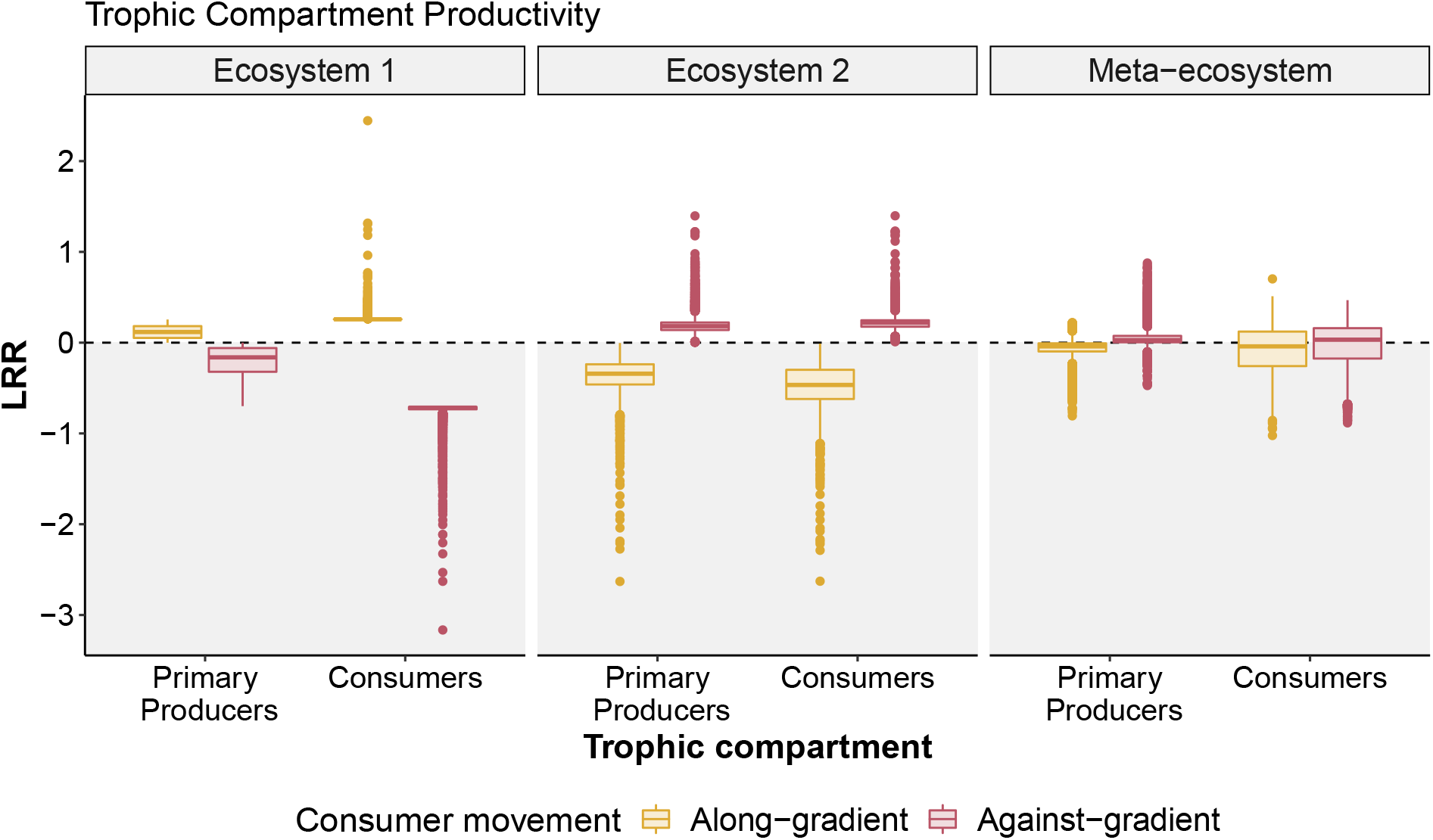
Changes to trophic compartment productivity when consumers move along-(yellow) or against-gradient (red) from ecosystem 1 to ecosystem 2. Ecosystem 1 sees a marginal increase in secondary productivity following along-gradient consumer movement (yellow) compared to the control scenario, whereas ecosystem 2 shows strong corresponding reductions for both primary and secondary productivity. However, the meta-ecosystem remains at productivity levels comparable to those of the control scenario. For against-gradient consumer movement (red), the situation is reversed with ecosystem 2 showing higher median levels of primary and secondary productivity compared to the control scenario, and ecosystem 1 having both functions reduced. Nevertheless, the meta-ecosystem shows only a marginal increase in secondary productivity and no change in primary productivity. All specifications as in Figure 2.

### 3.1 Consumers moving against the resource availability gradient

At the meta-ecosystem scale, against-gradient consumer movement does not appear to influence either nutrient flux (median *LRR*_*P*_ = 0.01, range: 0.00–0.90; *LRR*_*C*_ = 0.07, range: −0.86–0.49; Figure 3) or productivity (*LRR*_*P*_ = 0.03, range: −0.47–0.88; *LRR*_*C*_ = 0.03, range: −0.88–0.47; Figure 4), compared to the control scenario of homogeneous nutrient availability. Likewise, nutrient stock and trophic compartment biomass appear unchanged in the meta-ecosystem when movement happens against the nutrient availability gradient (*LRR*_*N*_ = 0.03, range: −0.67–0.25; *LRR*_*P*_ = 0.01, range: 0.00–0.90; *LRR*_*C*_ = 0.08, range: −1.16–0.59; Figure 2).

At the local ecosystem scale, we observe a different situation. Against-gradient consumer movement leads to the recipient ecosystem having higher primary (*LRR*_*P*_ = 0.18, range: 0.00–1.40) and secondary productivity (*LRR*_*C*_ = 0.22, range: 0.01–1.40) than in the control scenario (central panel, Figure 4). Consumer nutrient flux in the recipient ecosystem increases when consumers move against gradient (*LRR*_*C*_ = 0.17, range: −0.80–0.92), whereas primary producers nutrient flux appears unchanged (*LRR*_*P*_ = 0.04, range: 0.00–1.46) compared to the control (Figure 3). In turn, the donor ecosystem shows reduced values for consumer productivity (*LRR*_*C*_ = −0.72, range: −3.16– −0.70; Figure 4), nutrient flux (*LRR*_*C*_ = −0.72, range: −3.16– −0.70; Figure 3), and biomass accumulation (*LRR*_*C*_ = −0.72, range: −3.16– −0.70; Figure 2). Similarly, we observe reduced primary productivity (*LRR*_*P*_ = −0.16, range: −0.70–0.00) and nutrient stock accumulation (*LRR*_*N*_ = −0.16, range: −0.70–0.00) in the donor ecosystem compared to the control (Figures 2 and 4). Furthermore, our GSA reveals that consumer assimilation efficiency (*e*_2_) and consumer biomass loss rate (*d*_2_) in the recipient ecosystem are key determinants of the primary producers biomass in the donor ecosystem (see parameters *e*_2_ and *d*_2_ in the bottom row of Figure A.7). However, as noted above, the increase in local functioning of the recipient ecosystem appears to balance these reductions. Hence, while against-gradient consumer movement increases heterogeneity among local ecosystems, this difference is less pronounced at the meta-ecosystem level relative to the control scenario. Finally, we see limited to no change in biomass distribution following against-gradient consumer movement, for most parameter sets (see Appendix A.2 for details). Therefore, subsidy-induced shifts in the shape of local and meta-ecosystem biomass distribution have little impact on the patterns of local and meta-ecosystem functioning described above.

### 3.2 Consumers moving along the resource availability gradient

Along-gradient consumer movement does not appear to elicit marked changes compared to the control scenario at the meta-ecosystem scale (Figures 2 to 4). All three meta-ecosystem functions do not appear to vary with consumer movement (median *LRR* ∼ 0; Figures 2 to 4). The sole exceptions are changes in consumer biomass (*LRR*_*C*_ = −0.09, range: −1.08–0.57; Figure 2) and consumer-mediated nutrient flux (*LRR*_*C*_ = −0.09, range: −1.06–0.57; Figure 3). At the local ecosystem scale, however, along-gradient consumer movement has effects that are somewhat opposite to those of against-gradient movement. Along-gradient consumer movement leads to an increase in secondary productivity (*LRR*_*C*_ = 0.26, range: 0.26–2.45; Figure 4) and consumer nutrient flux (*LRR*_*C*_ = 0.26, range: 0.25–2.45; Figure 3) in the donor ecosystem, compared with the control. As well, we see a reduction of productivity and nutrient flux for both consumers (productivity *LRR*_*C*_ = −0.47, range: −2.63– −0.01; nutrient flux *LRR*_*C*_ = −0.27, range: −1.35–0.31) and primary producers (productivity *LRR*_*P*_ = −0.34, range: −2.63–0.00; nutrient flux *LRR*_*P*_ = −0.12, range: −2.72–0.00) in the recipient ecosystem compared to the control (Figures 3 and 4).Finally, as with against-gradient consumer movement, we observe limited to no change in biomass distribution following along-gradient movement, with little impact on the ecosystem functions patterns described above (see Appendix A.2 for details). In this scenario, parameters related to the functional response of consumers (*e*_1_) and of primary producers (*u*_1_) in the donor ecosystem appear to be the most influential parameters in shaping local consumer biomass and inorganic nutrient stock, respectively, according to our GSA (see middle row; Figure A.7). However, their influence does not cross ecosystem border to exert indirect effects on the biomass dynamics of the recipient ecosystem, which are molded by the locally important *u*_2_, *a*_2_, and *e*_2_ parameters for inorganic nutrients, primary producers, and consumers, respectively (middle row; Figure A.7).

## 4 Discussion

Organismal movement bridges ecosystem functions and processes across space and time, mediating exchanges of disparate currencies (Marleau, Peller, et al. 2020; Little et al. 2022) and connecting ecosystems together into meta-ecosystems (Massol, Gravel, et al. 2011; Massol, Altermatt, et al. 2017). Meta-ecosystem models generally account for organismal movement as a diffusive process, such as abiotic nutrient flows across ecosystems (but see Leroux and Loreau 2012; Häussler, Ryser, and Brose 2021). Consumer movement, however, is a multi-faceted process (Nathan et al. 2008; Earl and Zollner 2017) and a diffusion-like approach may not capture the variety of organism movement types observed in nature (Massol, Altermatt, et al. 2017; Gounand, Harvey, et al. 2018; McInturf et al. 2019). We integrate diffusive and non-diffusive (i.e., along- and against-gradient) movement into a novel meta-ecosystem model and investigate how different consumer movement types may influence ecosystems functions. Our results show that diffusive and non-diffusive movement of organisms have different, pervasive, direct and indirect effects on biomass and stock accumulation, productivity, and nutrient flux at local and meta-ecosystem extents. Analysis of our model highlights how the widespread assumption of diffusive consumer movement captures only a small fraction of local and meta-ecosystem dynamics. Further, our model provides testable predictions for consumer movement scenarios commonly observed in nature and can be a flexible tool to help bridge empirical and theoretical meta-ecology. We discuss ways to further expand our model and test predictions arising from it in both controlled and real-world scenarios.

Irrespective of gradients, consumers moving in meta-ecosystems directly influence consumer-mediated biomass accumulation (Figure 2), nutrient flux (Figure 3), and productivity (Figure 4) in both donor and recipient ecosystems. Key differences emerge, however, between movement that is against- or along-gradient (i.e., diffusion). Against-gradient, non-diffusive consumer movement results in stark differences between the donor and recipient ecosystems, with the latter showing increased stock and biomass accumulation, nutrient flux, and productivity (Figures 2 to 4). These differences balance out at the meta-ecosystem extent, resulting in less pronounced differences with the control or along-gradient scenarios (right-most panels, Figures 2 to 4). A real-world example of these dynamics may come from central-place foragers, such as beavers (*Castor* spp.) and some baleen whales (e.g., *Megaptera novaeangliae*). Through transport of biomass and nutrients from peripheral to central ecosystems, central-place forages have pervasive effects on the functions of local and meta-ecosystems that can last for decades after their disappearance, from increased nutrient availability and mineralization to higher nutrient cycling (see reviews in Rosell et al. 2005; Roman, Estes, et al. 2014). Conversely, along-gradient, diffusion-like consumer movement leads to the recipient ecosystem acting as a consumer biomass sink, in turn leading to locally reduced consumer-mediated productivity and nutrient flux (yellow box plots, Figures 2 to 4). Yet, at the meta-ecosystem scale, we do not see marked changes in the levels of biomass accumulation (Figure 2), nutrient flux (Figure 3), and trophic productivity (Figure 4) compared to those of an homogeneous landscape. Importantly, while consumer movement impacts biomass distribution at local and meta-ecosystem extents, our results are relatively robust to shifts in the distribution of biomass in the donor, recipient, or meta-ecosystem—i.e., from bottom-to top-heavy biomass pyramids (McCauley et al. 2018, see Appendix A.2 for details). Thus, a first key insight arising from our model is that, while the effects of along-gradient consumer movement align with well-known meta-ecosystem source-sink dynamics (Gravel, Guichard, et al. 2010), against-gradient consumer movement appears to increase the patchiness of nutrient concentration and availability over landscapes. This “resource pooling” effect mediated by against-gradient consumer movement may be instrumental in the emergence of temporal hot-spots of nutrient availability over the landscape (sensu Bernhardt et al. 2017), or in facilitating habitat turnover dynamics (McNaughton 1990).

Consumer movement across ecosystems can also have indirect effects on the dynamics of local and meta-ecosystems, with again key differences between against- and along-gradient movement. Moving consumers transfer foraging pressure from autotrophs in the donor to autotrophs in the recipient ecosystem, leading to the emergence of a spatial trophic cascade (sensu Knight et al. 2005; Monk and Schmitz 2022). When consumers move against-gradient, this cascade stimulates primary productivity (Figure 4) and nutrient flux (Figure 3) while keeping local autotroph biomass low in the recipient ecosystem. Conversely, the donor ecosystem sees a reduction of nutrient stocks (Figure 2) and productivity (Figure 4), which may relate to either reduced nutrient recycling or autotroph uptake, or both (Figure A.7). As a result, at the regional, meta-ecosystem scale, against-gradient consumer immigration in the recipient ecosystem maintains and exacerbates landscape heterogeneity—meaning, for instance, that dark green polygons in Figure 1a would become darker and light green polygons would get paler. Conversely, autotroph-mediated nutrient flux and productivity both decrease following along-gradient consumer movement, reducing nutrient accumulation in the recipient ecosystem (yellow box plots, Figures 2 to 4)—effects that are consistent with source-sink dynamics at the meta-ecosystem scale (Gravel, Guichard, et al. 2010). Furthermore, indirect effects of consumer movement can arise from the inherent redistribution of biomass in the meta-ecosystem it causes, altering or even inverting biomass pyramids at local and regional scales (Figures A.3 and A.5 in Appendix A.3; McCauley et al. 2018). Indeed, allochthonous subsidies can lead to shifts from classic to inverted biomass pyramids (Trebilco, Baum, et al. 2013). In our analyses inverted biomass pyramids arise from a limited number of parameter sets—≈4 % of stable parameter sets, across both experimental scenarios—and our analyses are robust to these changes (see Figures A.4 and A.6 in Appendix A.3). Nonetheless, there appears to be a region of parameter space where ecosystems with top-heavy biomass distribution can persist over time—possibly through the influence of consumer movement. In turn, this opens further research opportunities to investigate whether consumer movement—regardless of its type (Figures A.4 and A.6)—can act as a stabilizing force for ecosystems where biomass concentrates at the top of the food chain (e.g., in marine ecosystems; Trebilco, Baum, et al. 2013; Trebilco, Dulvy, et al. 2016).

Simple and diffusion-based consumer movement models abound in meta-ecosystem ecology and we surmise that one of the primary reasons for this is that representing movement in different ways leads to challenges in mathematical model analysis (Massol, Altermatt, et al. 2017). As such, a major contribution of our work is the development of a modeling framework to consider more diverse types of consumer movement while maintaining some level of analytical tractability. We achieve this by integrating two novel approaches into a classic meta-ecosystem model. First, we model the movement of consumers in our meta-ecosystem using a dynamical variable, 𝒬, that lets us explicitly account for spatial heterogeneity in ecosystem components (Figure 1a, c; Weisser and Hassell 1996). In turn, this allows us to separate local and regional dynamics, and to quantify the influence of consumer movement across spatial scales. Organisms can often make active decisions on when and where to move in a landscape (Nathan et al. 2008), and these vary with spatial scales (Johnson 1980). Stimuli and information collected from both the surrounding environment and the internal state of a consumer can further shape these decisions (Earl and Zollner 2017; Subalusky and Post 2019; McInturf et al. 2019). The dispersers’ pool we introduce may allow future meta-ecosystem models to better account for these contextual drivers of consumer movement—especially, the matrix between ecosystems—which are increasingly easier to quantify empirically (Ellis-Soto et al. 2021). For instance, while 𝒬 in our model acts as a transit state, future models could incorporate elements in eq. (1g) to account for partial or bidirectional movement (e.g., by introducing emigration of *C*_2_ towards 𝒬 Peller, Guichard, and Altermatt 2023), qualitative or quantitative ecosystem heterogeneity (e.g., stoichiometric patch quality; Leroux, Vander Wal, et al. 2017; Subalusky and Post 2019), the presence of competitors or predators (McInturf et al. 2019), or the configuration of the landscape the movement takes place in (Crespo-Pérez et al. 2011; McLeod and Leroux 2021).

The second novel approach we integrate in our model is the use of time scale separation (TSS; Otto and Day 2011) to tease apart concurrent processes in the meta-ecosystem. Time is a key variable for meta-ecosystem dynamics, defining and influencing both ephemeral (e.g., nutrient availability; Bernhardt et al. 2017) and long-term processes (e.g., ecosystem development; Menge, Hedin, and Pacala 2012). TSS has a long history of helping to study ecosystem processes that occur over different and overlapping time scales. For example, TSS helped develop accurate predictions of the dynamics of spruce budworm (*Choristoneura fumiferana*) outbreaks in the boreal forests of North America (Ludwig, Jones, and Holling 1978). TSS also helped shed light on the emergence of autotroph plasticity in nutrient uptake and limitation, showing that competing nutrient uptake strategies of plants vary in their adaptive value over time with environmental heterogeneity and can influence plant community dynamics and nutrient cycling (Menge, Pacala, and Hedin 2009; Menge, Hedin, and Pacala 2012). However, to our knowledge, our study is the first to apply TSS to a two-patch meta-ecosystem model to tease apart the effects of ecosystem processes at local and regional spatial extents. We can envision fruitful applications of the combined approach involving the dispersers’ pool and TSS in future meta-ecosystem ecology research. For instance, our assumption of *fast* consumer movement and *slow* biomass production may not hold in ecosystems other than terrestrial ones. Our proposed framework may help investigate consumer-mediated meta-ecosystem effects in systems where, for example, autotroph biomass production is much faster than consumer movement. As well, while most meta-ecosystem models focus on two-patch systems, real-world meta-ecosystem comprise several diverse environments. Moving beyond the simple formulation of 𝒬 that we propose could be instrumental to advance research on multi-patch, lattice, or even spatially explicit meta-ecosystem models.

Meta-ecosystem theory—and, indeed, meta-ecology at large—has historically focused on local and meta-ecosystem stability as independent variables of interest (Gravel, Guichard, et al. 2010; Marleau, Guichard, Mallard, et al. 2010; Marleau, Guichard, and Loreau 2014; Marleau, Peller, et al. 2020). However, ecosystem functions are diverse and encompass a broad range of processes that enable and complement ecosystem stability, because they feed back on each other. The modeling framework we propose allows us to pivot from just a focus on stability, to consider how feedbacks among local- and meta-scales arise from consumer movement that influences some of these ecosystem functions—namely, primary and secondary productivity and nutrient flux (sensu Peller, Guichard, and Altermatt 2023). Crucially, these ecosystem functions are commonly measured in empirical studies (Garland et al. 2021). For instance, measuring gross primary production and respiration rates of freshwater autotrophs was instrumental in disentangling the effects of subsidies on ecosystem functions mediated by different types of consumer movement in the Mara River, Kenya (Subalusky, Dutton, Njoroge, et al. 2018). As well, data on organic matter flow and secondary productivity from the Horonai Stream, Japan, helped show that consumermediated subsidies may elicit opposite effects on ecosystem functions at different temporal scales (Marcarelli et al. 2020). We hope that, by allowing integration of widespread measurements of ecosystem functions in mathematical models, our approach will foster the development of feedbacks between empirical and theoretical meta-ecology—with the potential for real-world applications. For instance, how might the removal or alteration of organismal movement pathways (e.g., through road or hydro-electric dam construction; Tucker et al. 2018) impact ecosystem productivity and nutrient flux at local and regional extents in, for example, the meta-ecosystem formed by salmon-spawning rivers and tropical boreal forests of the North American Pacific North-West. Or, in the wake of the COVID-19 pandemic, how widespread, rapid, and long-term changes in humankind’s habitat use patterns (e.g., the COVID-19 Anthropause; sensu Rutz et al. 2020) may in turn vary consumer movement pathways and the connections among, e.g., agricultural and forested areas in the Central European landscape mosaic (Abbas et al. 2012).

In our model, we focus on reproducing key dynamics of a complex phenomenon (i.e., movement of medium-to-large land mammals; Nathan et al. 2008) and make several simplifying assumptions. While open at the basal level, our model does not feature ecosystem flows other than apical consumer movement (Figure 1c), complementing and expanding earlier studies of basal, diffusive flows (Gravel, Guichard, et al. 2010; Gravel, Mouquet, et al. 2010; Marleau, Guichard, Mallard, et al. 2010) and of consumer influence on ecosystem dynamics (Leroux and Loreau 2010; Leroux, Hawlena, and Schmitz 2012; Leroux and Schmitz 2015). We see developing and analyzing models that include both diffusive and non-diffusive flows as a natural extension of our work. Our model further assumes one-way consumer movement. Real-world consumers rarely move so, and frequently travel back and forth between habitats (Gounand, Harvey, et al. 2018). The decision to move itself involves complex trade-offs influenced by both internal and external stimuli (Nathan et al. 2008; Little et al. 2022). Integrating concepts and tools from landscape ecology, where space is often treated explicitly, may help better account for the spatial relationship and configuration of the matrix-ecosystem mosaic (sensu Castilla et al. 2009; Wu 2013) and inform the formulation of 𝒬. For example, mapping patch compartments into a spatially explicitly cellular automata model (e.g., Crespo-Pérez et al. 2011) or using friction values in a spatial model to parameterize matrix portions with different permeability (e.g., Coulon et al. 2015) would be useful to test how landscape features might influence consumer movement. Further, deeper integration of landscape ecology and emerging information ecology ideas (Marleau, Peller, et al. 2020; Little et al. 2022) in our model may help account for human modifications of movement pathways. Habitat fragmentation (Haddad et al. 2015), sensory pollution (Sanders et al. 2021), and movement barriers (Tucker et al. 2018) can all potentially influence the trajectory and scale of organismal movement. Our model’s ability to account for these modifications using a dynamic variable (𝒬), while maintaining mathematical tractability, could help develop new hypotheses and predictions of their effects on both organismal movement and ecosystem functions. Field-based, multi-scale studies of meta-ecosystems and the flows that connect them—e.g., in the Pacific West (Menge, Gouhier, et al. 2015; Menge, Caselle, et al. 2019) or the Galápagos Islands (Witman, Brandt, and Smith 2010)—would be ideal candidates to explore the role of consumers-mediated subsidies in meta-ecosystems. In terrestrial ecosystems, extant herbivore exclusion experiments appear ideally suited as starting point to develop comparable studies spanning multiple spatio-temporal scales.

Meta-ecosystem ecology encourages and challenges researchers to expand their focus and look at general, emerging spatial properties of ecosystems. Here, we develop a novel, flexible approach to investigate the influence of different types of consumer movement in meta-ecosystems. Key results of our model include that (i) different types of consumer movement have different, at times opposite, influences on local and meta-ecosystem stocks, nutrient flux, and productivity; (ii) these influences can be either direct, developing within the same trophic compartment across ecosystem borders, or may be indirect and affect other compartments in the local food web; (iii) partitioning local and meta-ecosystem responses to processes that span spatio-temporal scales, like consumer movement, is key in unveiling feedbacks among local ecosystems that may influence local and meta-ecosystem functions and dynamics; and (iv) consumer movement, irrespective of its type, may act as a stabilizing force for ecosystems where biomass concentrates at the top of the food chain. Our results heed the call for integrating more complex representations of organismal movement in meta-ecosystem models (Massol, Altermatt, et al. 2017; Peller, Guichard, and Altermatt 2023) and show that current meta-ecosystem models based on the mathematically convenient assumption of diffusive, along-gradient consumer movement predict only a subset of dynamics that may be expected in meta-ecosystems. As humankind adopts new actions and strategies to mitigate the effects of anthropogenic environmental change, moving beyond long-established assumptions and accounting for the myriad moving pieces that shape ecosystem functioning has never been more pressing.

## 5 Acknowledgments

We thank J. Balluffi-Fry, I. C. Richmond, R. Buchkowski, and J. Monk for helpful comments during model development and on earlier drafts of this manuscript. We thank A. McLeod for pointing us to the Weisser and Hassell (1996) paper, and A. McLeod and R. Buchkowski for their help coding the stability analyses. We thank the Ecosystem Ecology Lab at Memorial University of Newfoundland for helpful conversations and support throughout this project. This research was funded by the Government of Newfoundland and Labrador Centre for Forest Science and Innovation to SJL (grant #221274), YW (grant #221273), and EVW (grant #221275), Government of Newfoundland and Labrador Innovate NL Leverage R&D to EVW & SJL (grant #5404.1884.102) and Ignite R&D to SJL (grant #5404.1696.101) programs, Mitacs Accelerate Graduate Research Internship program to YW, EVW, & SJL (grant #IT05904), the Canada Foundation for Innovation John R. Evans Leaders Fund to EVW & SJL (grant #35973), Natural Science and Engineering Research Council Discovery Grant to SJL (grant #RGPIN 435372-2013), Mitacs Globalink Research Award (#IT12980) and a Mitacs Research Training Award (#IT20836) to MR.

## 6 Data Availability

All code used in our analyses is available at: https://doi.org/10.6084/m9.figshare.16479933

## 7 Authorship Statement

MR and SJL conceptualized the idea; MR, SJL, OJS, EVW, YFW, and TRH designed the study; MR, SJL, and OJS developed the model; MR ran the analyses; MR, SJL, and OJS interpreted the results; MR led the writing of the manuscript; all authors contributed critically to the manuscript and approved the final version.

## A Appendices

Here, we provide additional details on the analyses of our model and on our results. In Appendix A.1, we report feasible and unfeasible equilibria of the model. In Appendix A.2, we describe in detail the results of our analyses on changes in the biomass distribution in our metaecosystem caused by consumer movement, and their effects on the top-down influence of consumer movement on local and meta-ecosystem functions. In Appendix A.3, we present additional figures for the untransformed results of our model iterations and for our biomass distribution analyses. Finally, in Appendix A.4, we provide the results of the stability analyses we performed on our model’s single feasible equilibrium.

### A.1 Model Equilibria

Here we report the feasible and unfeasible equilibria of our model (see text, eq. (1)). We derived these equilibria using Mathematica (v. 13.0, Wolfram Research Inc. 2020), and then transferred them to R (v. 4.2.0, R Core Team 2022) to run our simulations. As described in the main text, we first solve eq. (1g) to find the quasi-equilibrium 𝒬* (eq. (2)). We then substitute eq. (2) in eq. (1f) and solve for 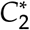. Through subsequent rounds of substitution, we solve for all other state variables. Finally, we substitute the solution for 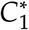 into eq. (2), to get the solution for 𝒬*. In the following formulae, we use these substitutions for clarity:

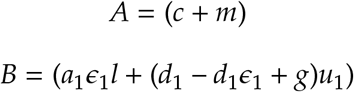

Feasible equilibria:

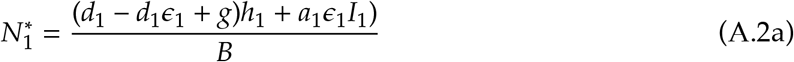

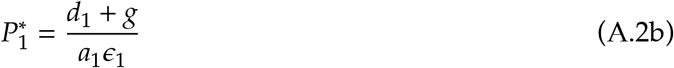

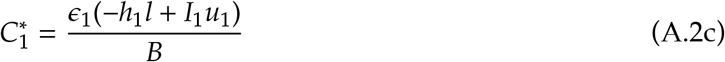

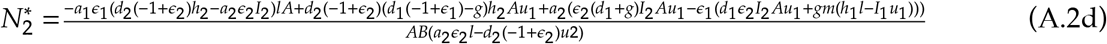

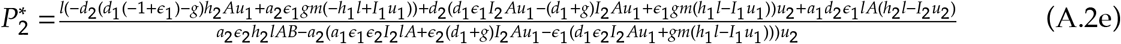

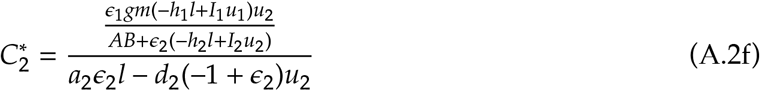

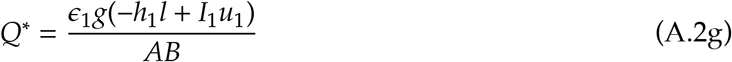

Other, unfeasible equilibria:

Case 1:

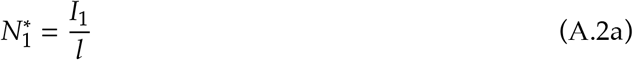

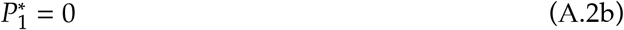

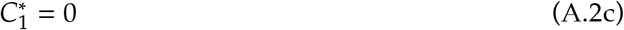

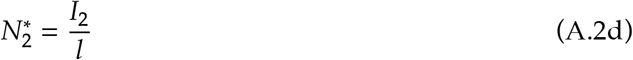

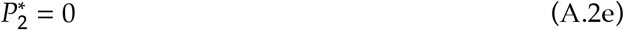

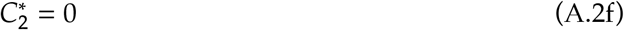

Case 2:

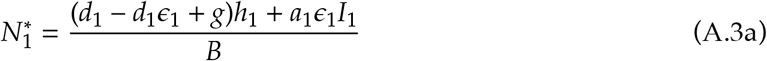

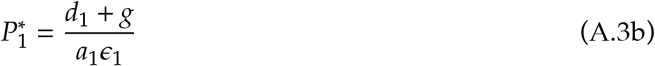

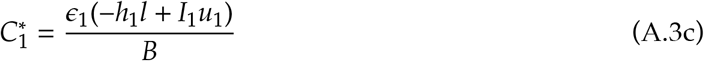

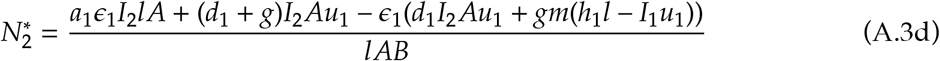

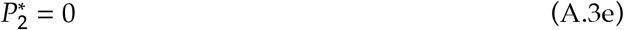

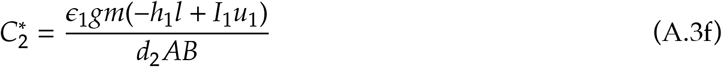

Case 3:

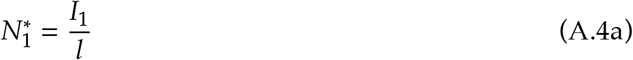

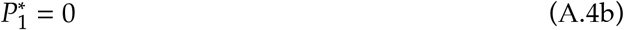

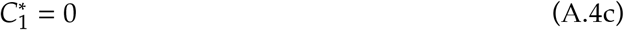

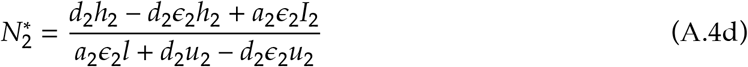

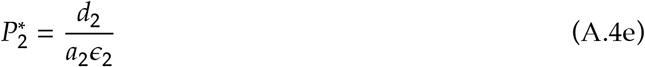

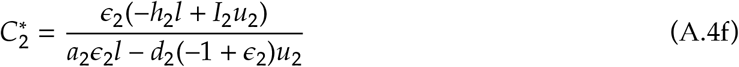

### A.2 Changes to meta-ecosystem biomass distribution with consumer movement

Consumers movement, whether along-or against-gradient, changes the distribution of biomass in the meta-ecosystem. Potentially, this can lead to changes in the shape of local and meta-ecosystem biomass pyramids from bottom-to top-heavy—as shown by recent empirical (Trebilco, Baum, et al. 2013; Trebilco, Dulvy, et al. 2016) and theoretical (McCauley et al. 2018) studies. We used the Consumer to Resource biomass ratio (*C*:*R*) to investigate any synergies between biomass pyramid shape and the different types of consumer movement under scrutiny in our work.

For both experimental movement scenario, along- and against-gradient consumer movement, most parameter sets produce a classic, bottom-heavy distribution of biomass at both local and regional spatial scales. That is, most parameter sets that produced stable model equilibria led to both the donor and recipient ecosystems, as well as the meta-ecosystem as a whole, to have a *C*:*R* ∈ [0, 1] (Figure A.3). A *C*:*R* within this range of values is generally interpreted as evidence of bottom-heavy biomass pyramids (e.g., Figure A.3; McCauley et al. 2018). In turn, this did not appear to qualitatively change our results (compare Figures 2 to 4 with panels (a) to (c) in Figure A.4, respectively). However, and perhaps more surprising, a few parameter sets produced stable equilibria where local and meta-ecosystems have inverted biomass pyramids when consumers move against-gradient (i.e., *C*:*R* ∈ (1, 10); e.g., Figure A.5). These are more peculiar, as inverted biomass pyramids are theoretically rare in nature but have been empirically documented in several ecosystems—mostly marine ones (e.g., Trebilco, Dulvy, et al. 2016). However, as we show the Supporting Code document in the Data and Code repository, our results remain unchanged when consumer movement leads to inverted biomass pyramids in the donor ecosystem 1, the recipient ecosystem 2, or the meta-ecosystem.

As such, we conclude that the results of our modeling effort are robust to consumer movement-induced changes in the distribution of biomass in the meta-ecosystem. However, we believe that further exploration of the subset of parameter space that leads to stable, biologically feasible equilibria for our model may be a fruitful line of research in the future. In particular, as consumer movement may be key in stabilizing meta-ecosystem where biomass concentrates at the top of the food chain in one or more ecosystems (Trebilco, Baum, et al. 2013).

### A.3 Additional Figures

Here, we present additional figures that provide more detail into our model’s result. Figure A.1 provides a visual aid to interpret the *log*_10_ Response Ratio results reported in Figures 2 to 3 and in Figures A.4 and A.6. In Figure A.2, the three panels report the untransformed values for the results shown in Figures 2 to 4. Figures A.3 and A.5 shows the shape of the biomass pyramid in either local ecosystem and in the meta-ecosystem when the donor ecosystem’s biomass distribution conforms to either a classic (Figure A.3) or an inverted (Figure A.5) pyramid. Figures A.4 and A.6 show the effects of along- and against-gradient consumer movement on the three ecosystem functions of interest, split by the shape of the biomass pyramids in the local and meta-ecosystem—bottom- and top-heavy, respectively. Finally, Figure A.7 shows the relative importance of each parameter in the model across the three consumer movement scenarios.

**Figure A.1:**
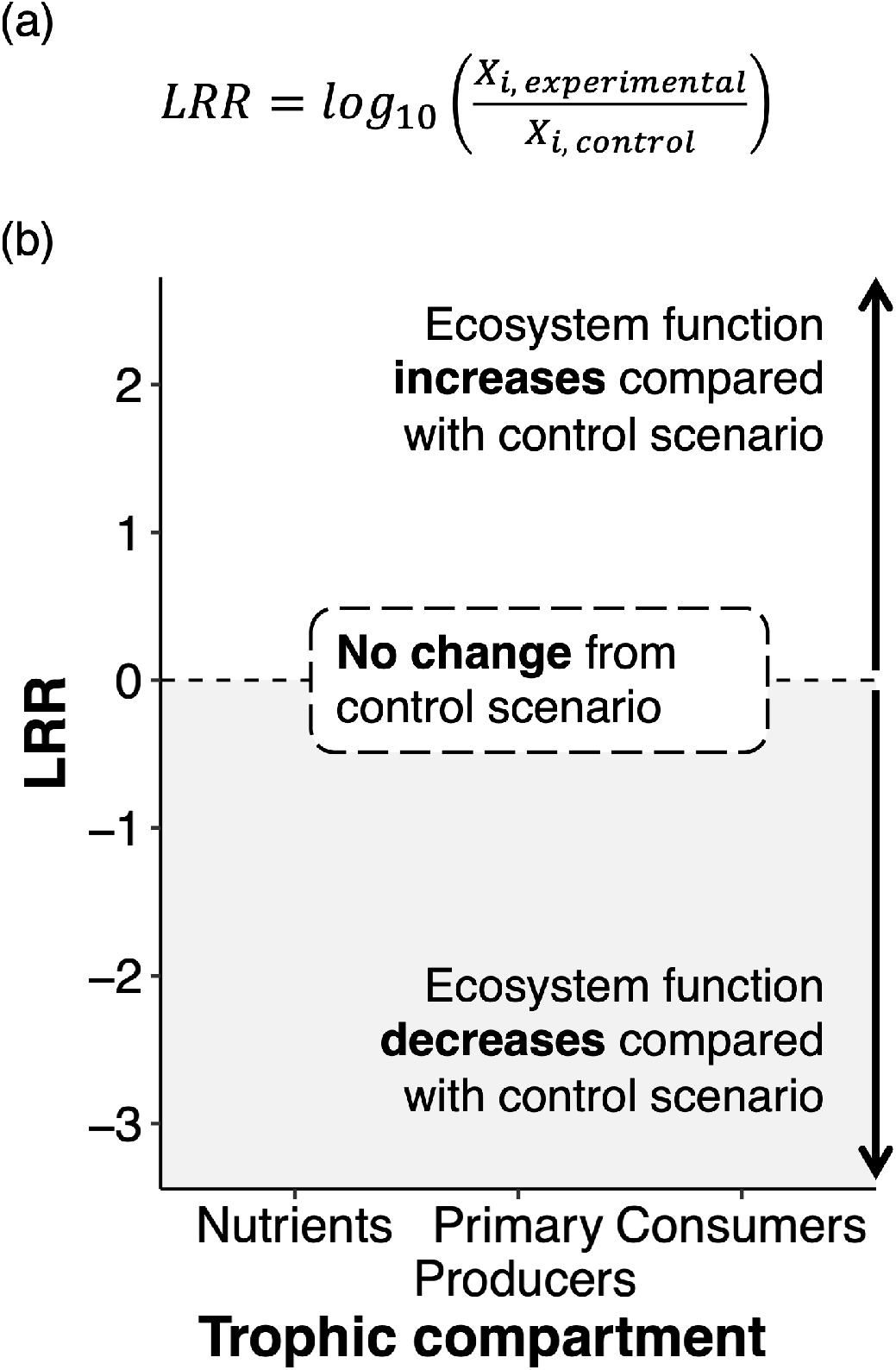
Visual aide to help interpret the results presented in Figures 2 to 4, as well as in Figures A.4 to A.6. **Panel (a)**: The *log*_1_0 Response Ratio (LRR) captures the change in the value of an ecosystem function of interest in one of the two movement scenario—along- and against-gradient—compared to the control, gradient-neutral movement scenario. In the equation, *X* represents one of biomass and stock accumulation, nutrient cycling, and trophic compartment productivity, and *i* ∈ [1, 2] represents either ecosystem in the model. “Experimental” is either the along-or the against-gradient movement scenario, whereas “control” stands for the gradient-neutral movement scenario. **Panel (b)**: Accordingly, in the Figures 2 to 4 and in Figures A.4 to A.6, the dashed black line at *LRR* = 0 represents no change in the ecosystem function of interest from the control scenario. When *LRR* > 0, the value of the ecosystem function of interest is higher than in the control scenario. Conversely, *LRR* < 0 points to the value of the ecosystem function of interest is lower than in the control scenario.

**Figure A.2:**
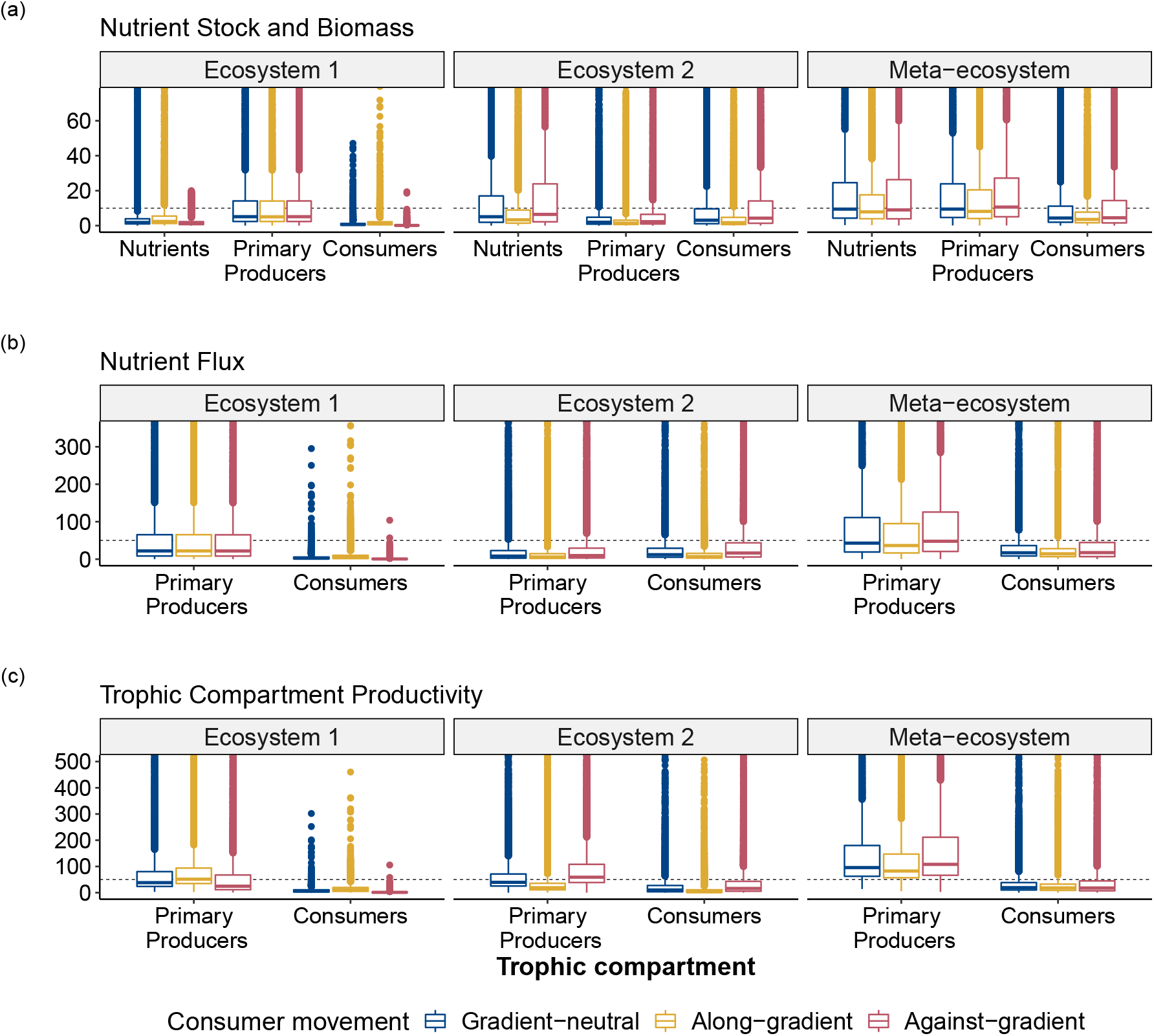
Change in (a) nutrient stock and biomass, (b) nutrient flux, and (c) trophic compartment productivity untransformed values, with different consumer movement types. Consumer move from ecosystem 1 to ecosystem 2, establishing a spatial trophic cascade. At the metaecosystem scale, function values appear higher for the against-gradient movement scenario (red) than for either the gradient-neutral (blue) or along-gradient (yellow) scenarios. Increased stock, nutrient flux, and productivity are likely a consequence of consumers movement from a less to a more nutrient-rich ecosystem exacerbating the spatial trophic cascade (compare left and central panels in (a)). Conversely, the lower functions values observed for the along-gradient scenario (yellow) likely point to the spatial trophic cascade enabling source-sink dynamics between local ecosystems. Blue box plots show function values for the control scenario of gradient-neutral consumer movement—i.e., the denominator in the *LRR* used in Figures 2 to 4. For each ecosystem function, thick lines inside the boxes represent median values, the lower (upper) hinge is the 25% (75%) quartile, and the lower (upper) whisker extends from the hinge to the smallest (largest) value no further than 1.5×interquartile range. Filled points beyond whiskers are outliers. Note the different scales of the ordinate. The dashed line is meant to help in interpreting the relative magnitudes of the function values plotted; it intercepts the y-axis at *y* = 10 for panel (a) and at *y* = 50 for panel (b) and panel (c).

**Figure A.3:**
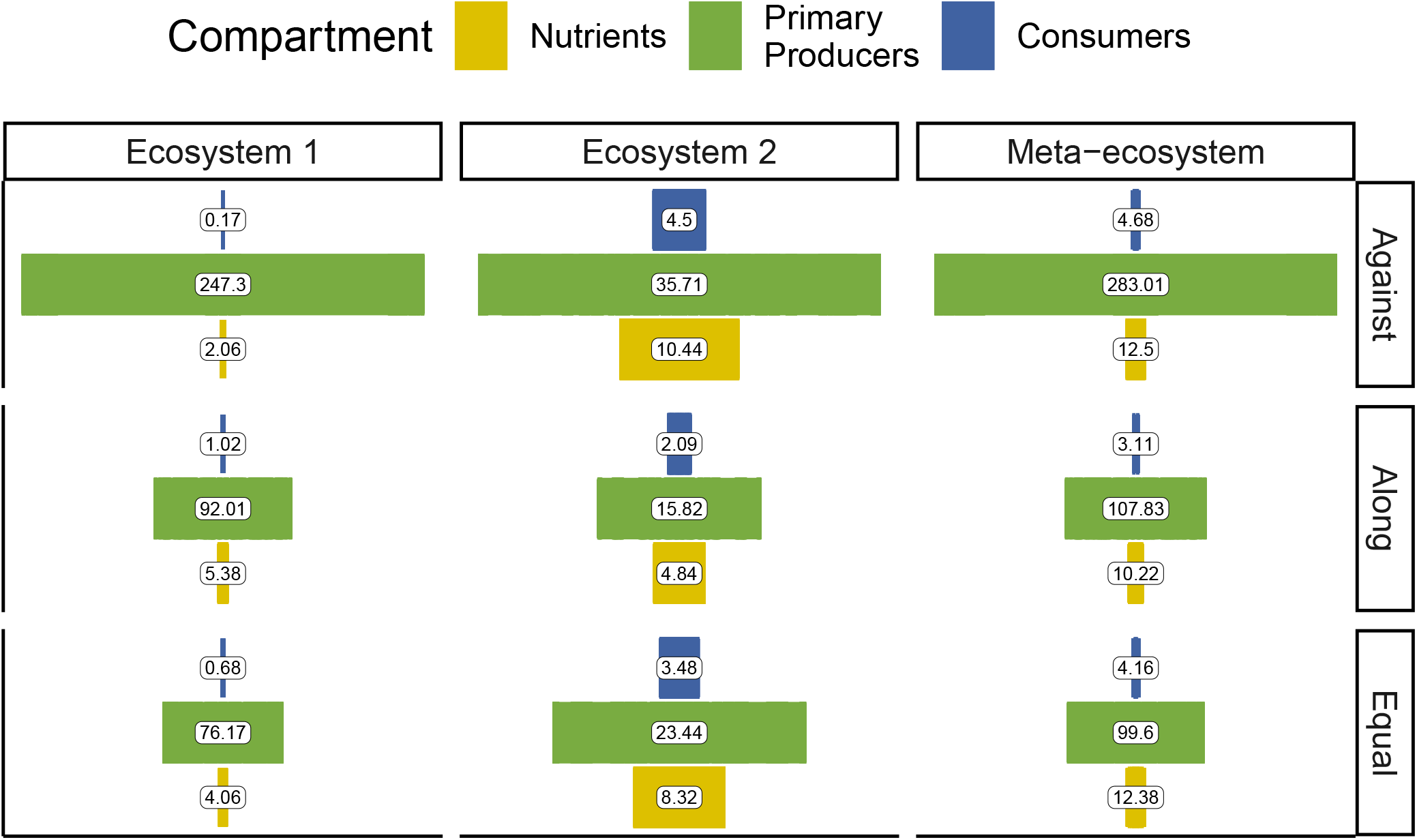
Visual representation of biomass distribution in the donor ecosystem 1, the recipient ecosystem 2, and the meta-ecosystem when ecosystem 1 has a classic, bottom-heavy biomass distribution (left column). When the donor ecosystem 1 conforms to a classic biomass pyramid, with a large base of primary producers (green) and small quantity of consumers (blue), both the recipient ecosystem 2 and the meta-ecosystem reflect a similar distribution of biomass. In turn, this has consequences for the effects of consumer movement in the system (Figure A.4). Note, however, the difference in the Consumer:Resource ratio between the donor ecosystem 1 and the recipient ecosystem 2. As well note that, in assessing the shape of biomass distribution, we exclude the inorganic Nutrient compartment (yellow). The labels inside the bars show the average biomass (g) for each ecosystem compartment.

**Figure A.4:**
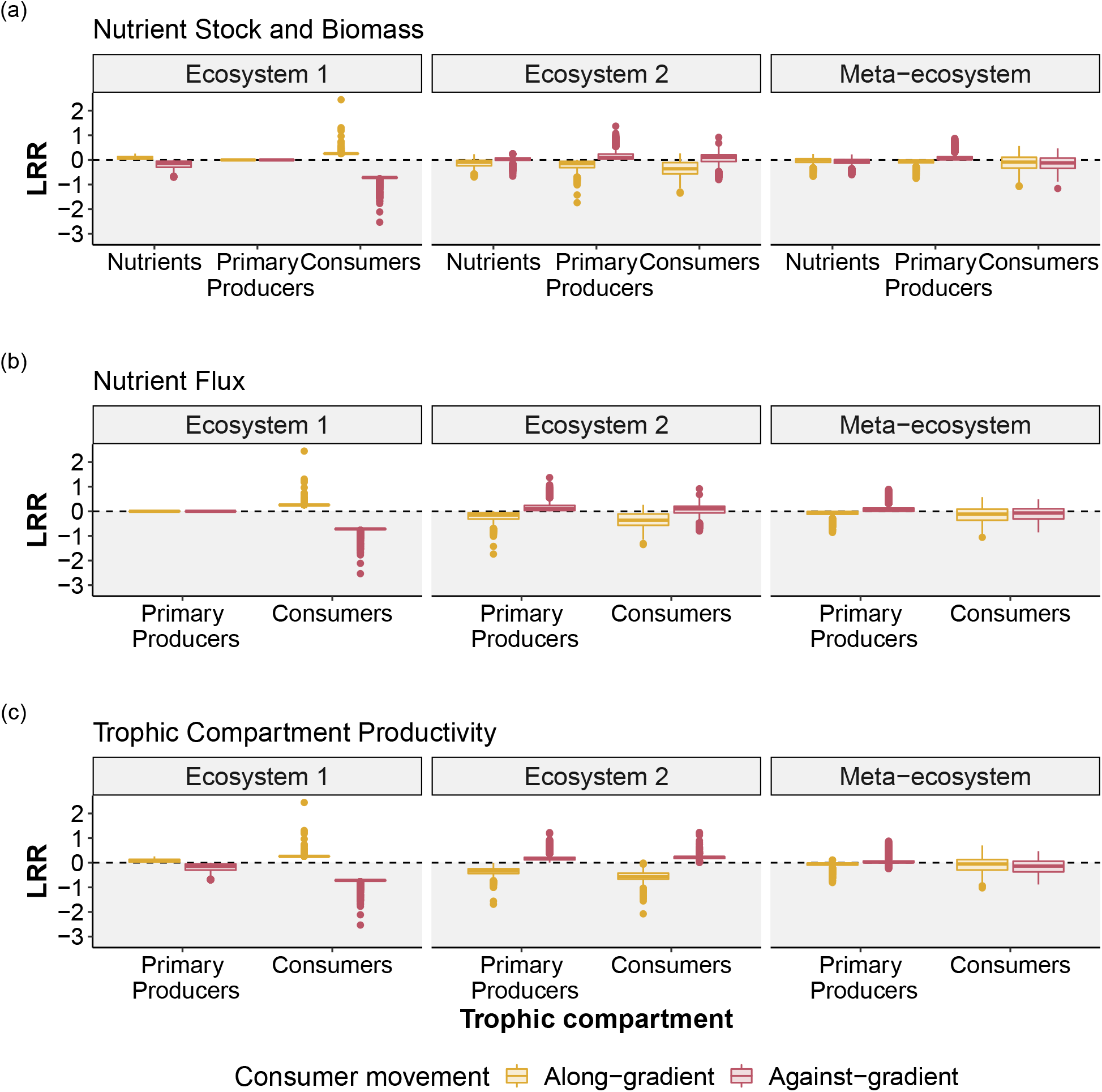
Here we show the effects of consumer movement on local and meta-ecosystem functions when the donor ecosystem has a classic, bottom-heavy biomass pyramid (i.e., *C*:*R* ∈ [0, 1]). Consumer movement from the donor ecosystem 1 to the recipient ecosystem 2 establishes a spatial trophic cascade. As in Figures 2 to 4, we observe a “resource pooling” effect when consumers move against-gradient of resource availability, so that resource-rich local ecosystems become increasingly richer. Along-gradient consumer movement instead leads to a homogenization of the resource distribution over the landscape. In both scenarios, local changes balance each other at the regional, meta-ecosystem scale. Note the *log*_10_ scale on the ordinate. All other specifications as in Figure 2.

**Figure A.5:**
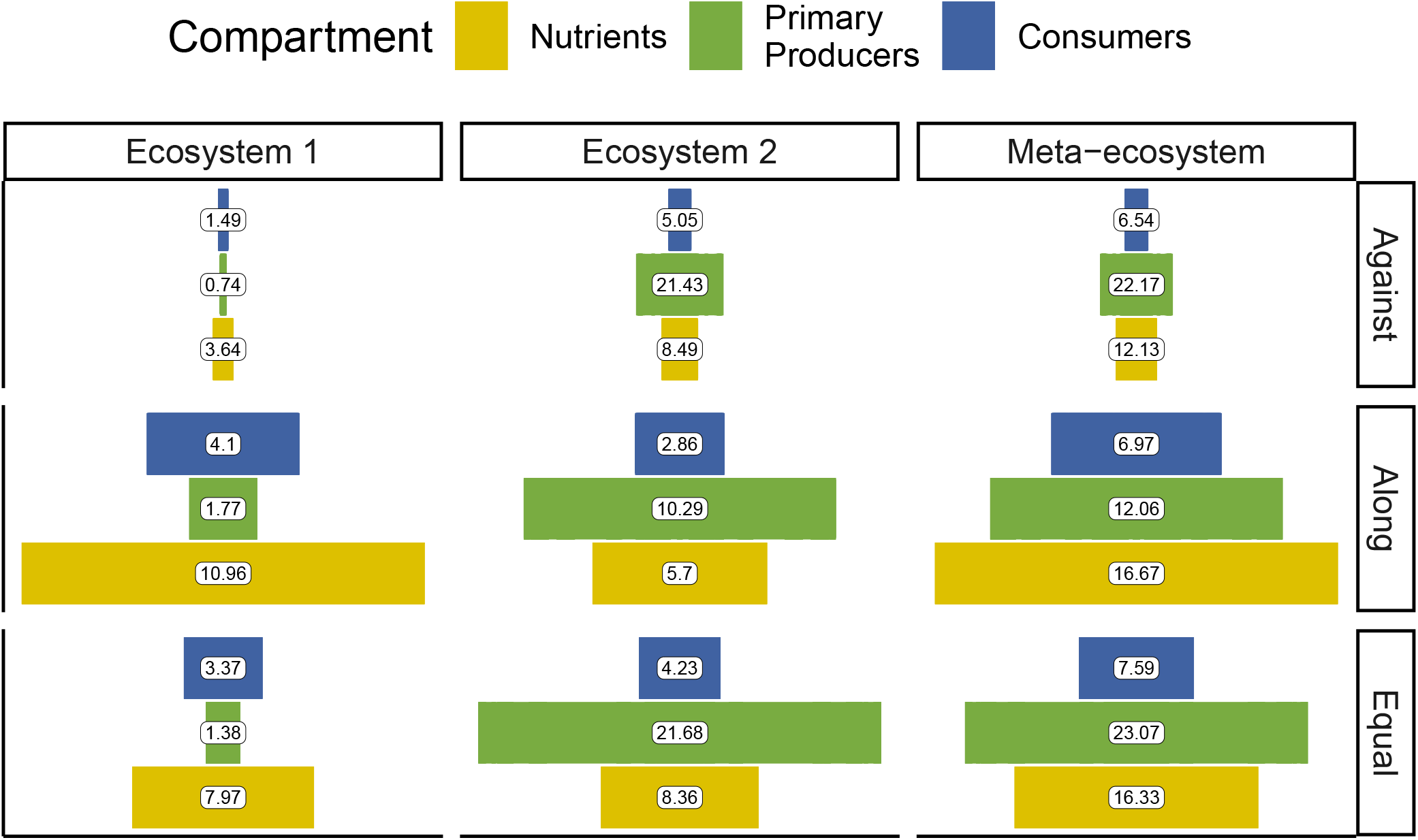
Biomass pyramids for the donor ecosystem 1, the recipient ecosystem 2, and the meta-ecosystem when the donor ecosystem 1 shows an inverted, top-heavy biomass distribution. This diagram captures those model runs where consumer biomass (blue) in the donor ecosystem 1 was much more abundant than primary producers biomass (green), across all three consumer movement scenarios. As the central and right-most column show, an inverted biomass pyramid in the donor ecosystem 1 does not result in a similar distribution of biomass in the recipient ecosystem 2 or in the meta-ecosystem. In turn, this influences the way consumer movement effects play out at local and regional scales (Figure A.6). All specifications as in Figure A.3. The labels inside the bars show the average biomass (g) for each ecosystem compartment.

**Figure A.6:**
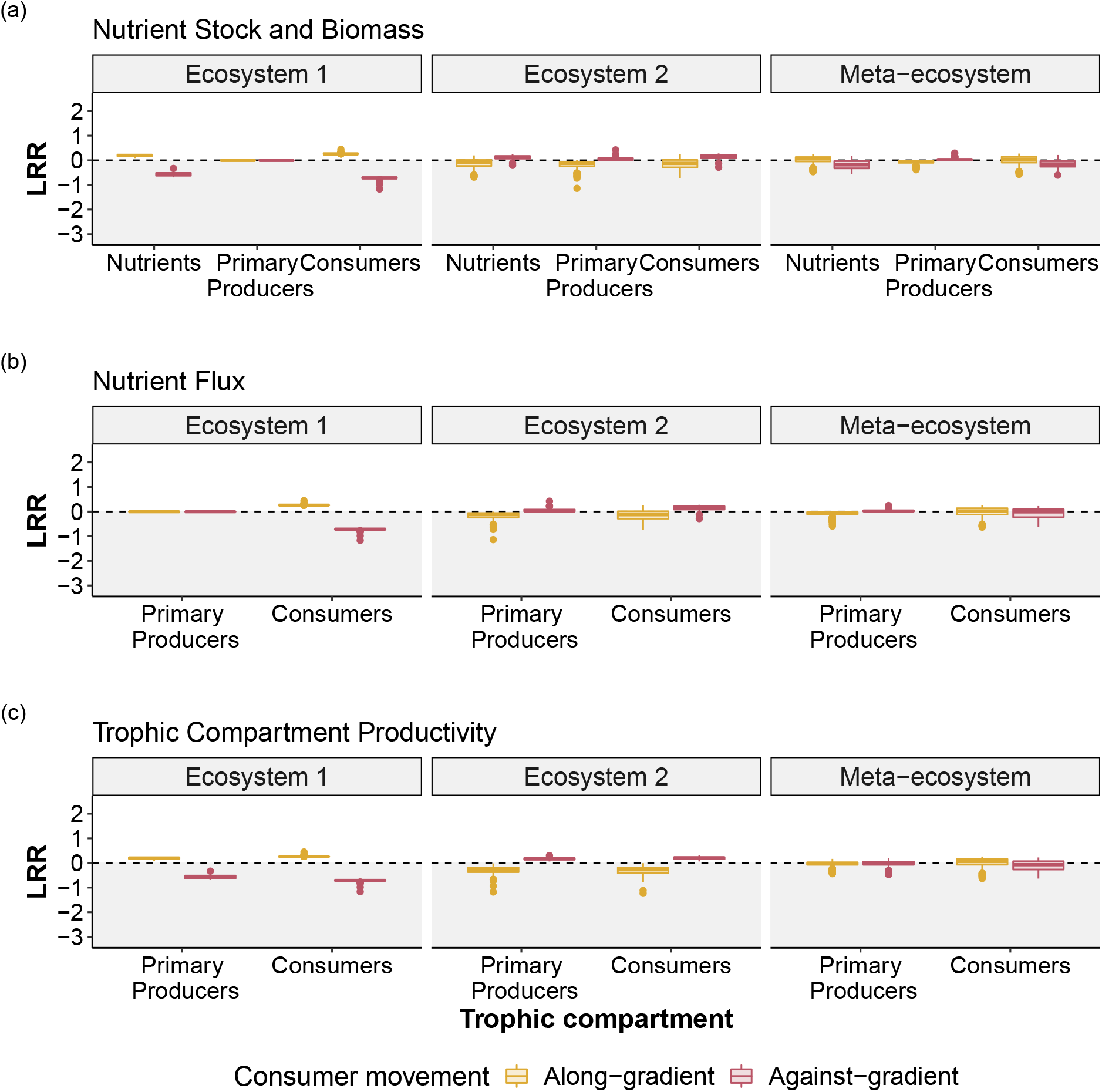
In contrast with Figure A.4, here the donor ecosystem has an inverted biomass pyramid (*C*:*R* ∈ (1, 10)). However, the effects of consumer movement on local and meta-ecosystem functions are qualitatively equivalent to those reported in Figures 2 to 4. Consumer movement from ecosystem 1, the donor, to ecosystem 2, the recipient, establishes a spatial trophic cascade between local ecosystems. Against-gradient consumer movement results in higher functioning in the recipient ecosystem 2, consistent with a “resource pooling” effect in resource-rich areas of the landscape (red box plots). Along-gradient consumer movement, conversely, leads to homogenization of the resource distribution over the landscape (yellow box plots). Notably, few parameter sets produce stable systems with inverted biomass pyramids. Thus, while consumer movement may act as a stabilizing force in these systems, this stabilization effect appears confined to a narrow region of the parameter space. In turn, this reinforces expectations of inverted biomass pyramids being relatively rare in nature. Note the *log*_10_ scale on the ordinate. All other specifications as in Figure 2.

**Figure A.7:**
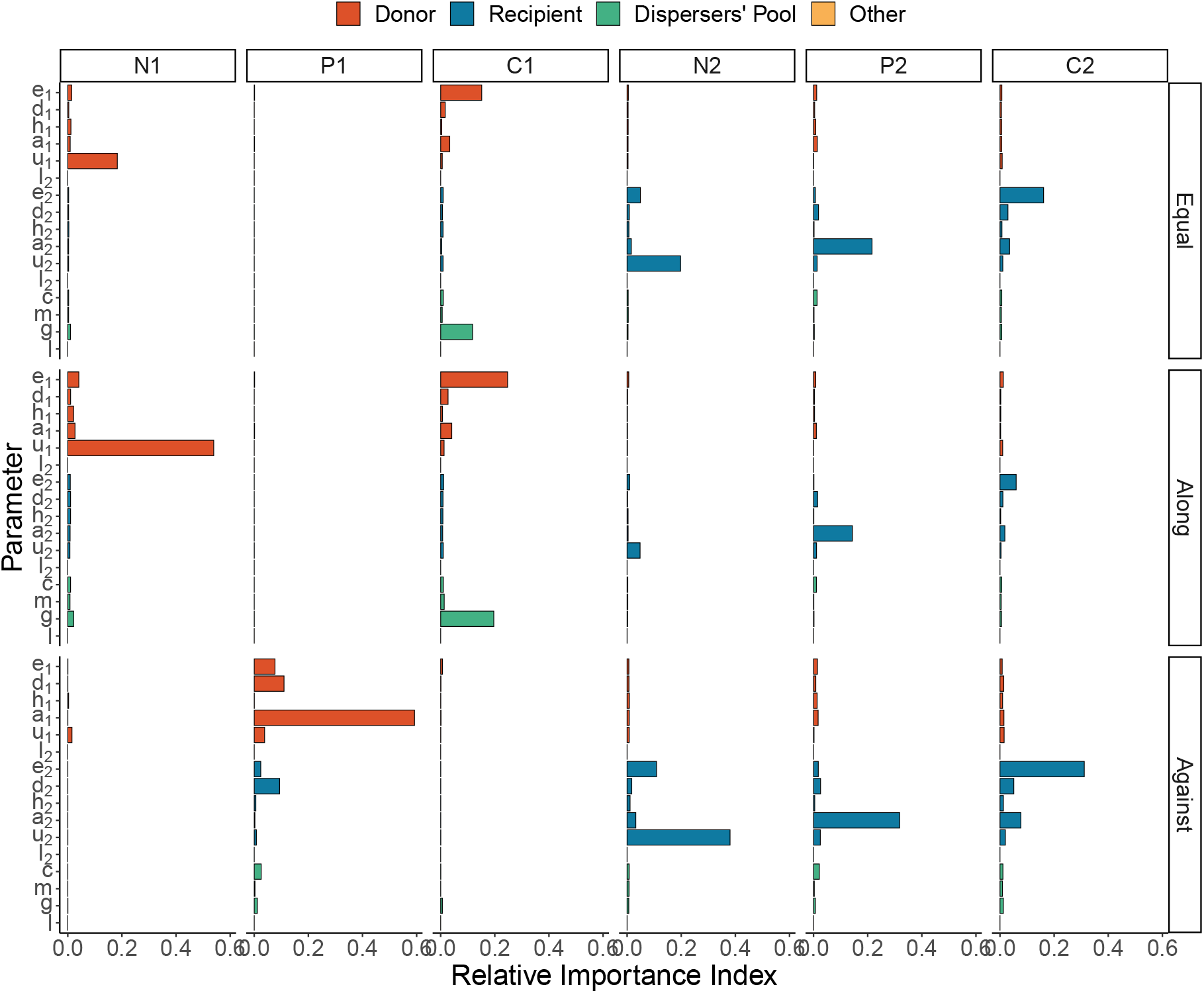
Relative importance of the parameters in the model, for each trophic compartment across consumer movement scenarios. **Top row**: for gradient-neutral movement (*I*_1_ = *I*_2_), the more important parameters in the donor ecosystem 1 are the primary producers’ uptake rate between *N*_1_ and *P*_1_ (*u*_1_) and the trophic efficiency of *C*_1_ (*e*_1_). Notably, in this ecosystem, we do not see evidence of indirect, top-down effects of consumers. Conversely, in the recipient ecosystem 2, we see evidence of an indirect effects as the second most important parameter for *N*_2_ is the consumers’ efficiency rate (*e*_2_), hinting at top-down control by *C*_2_ on nutrient dynamics. **Middle row**: when consumers move along-gradient, we see no evidence of indirect effects at play in either local ecosystem. All the more important parameters for each trophic compartment in either ecosystem pertain to the direct trophic interactions between them. **Bottom row**: when consumer move against-gradient, we see evidence of indirect, top-down effects within and across ecosystems. In the recipient ecosystem 2, the consumers’ efficiency (*e*_2_) appears to drive nutrient dynamics together with the primary producers’ uptake rate (*u*_2_). As well, and in contrast with what happens in the other movement scenario, functional response parameter of consumers in the recipient ecosystem 2 appear to influence the dynamics of primary producers in the donor ecosystem 1. Colors and grouping along the y-axis denote the meta-ecosystem component each parameter belongs to: the donor ecosystem 1 (red), the recipient ecosystem 2 (blue), or the dispersers’ pool (teal). Note the only parameter belonging to group “Other” (orange) is the leaching rate *l*, which has the same value across local ecosystems and in every consumer movement scenario.

### A.4 Model Stability Analyses Results

Table A.1 shows the results from the stability analyses of our model. We assessed the model stability by checking if the leading eigenvalue of the model’s Jacobian matrix (i.e., the eigenvalue with the largest real part; Otto and Day 2011) was positive or negative. Positive leading eigenvalue identify unstable equilibria, which we excluded from all analyses. We repeated this procedure for all parameter sets used in our analyses (n = 10 000) and for all three scenarios tested. For further details on our stability analyses, see the Supporting Code document in the data and code repository.

**Table A1:** Summary of stability analyses, showing the number of stable and unstable equilibria, and the percentage of unstable equilibria over the total number of parameter sets used (n = 10 000). A small portion of the parameter sets used in the analyses of our model produce unstable equilibria. This number varies among control and experimental scenarios. We excluded unstable parameter sets from all analyses. See the Supporting Code document in the online code repository for further details.

| Scenario | Stable | Unstable | % |
| --- | --- | --- | --- |
| Control | 9845 | 155 | 1.55 |
| Along-gradient consumer movement | 9676 | 324 | 3.24 |
| Against-gradient consumer movement | 9715 | 285 | 2.85 |

